# IQD1 involvement in hormonal signaling and general defense responses against *Botrytis cinerea*

**DOI:** 10.1101/2021.07.25.453677

**Authors:** Omer Barda, Maggie Levy

## Abstract

IQ Domain 1 (IQD1) is a novel calmodulin-binding protein in *A. thaliana*, which was found to be a positive regulator of glucosinolate (GS) accumulation and plant defense responses against insects. We demonstrate here that the IQD1 overexpressing line (*IQD1^OXP^*) is more resistant also to the necrotrophic fungus *Botrytis cinerea*, whereas an IQD1 knockout line (*iqd1-1*) is much more sensitive. Furthermore, we show that IQD1 is upregulated by Jasmonic acid (JA) and downregulated by Salicylic acid (SA). Comparison of whole transcriptome expression between *iqd1-1* and wild type revealed a substantial downregulation of genes involved in plant defense and hormone regulation. Further examination revealed a marked reduction of SA/JA signaling and increase in ethylene signaling genes in the *iqd1-1* line. Moreover, quantification of SA, JA and abscisic acids in *IQD1^OXP^* and *iqd1-1*lines compared to WT showed a significant reduction in endogenous JA levels in the knockout line simultaneously with increased SA levels. Epistasis relations between *IQD1^OXP^* and mutants defective in plant-hormone signaling indicated that IQD1 acts upstream or parallel to the hormonal pathways (JA/ET and SA) in defense response against *B. cinerea* and in regulating GS accumulation and it is dependent on JAR1 controlling indole glucosinolate accumulation. As a whole, our results suggest that IQD1 is an important defensive protein against *Botrytis cinerea* in *A. thaliana* and is integrated into several important pathways such as plant microbe perception and hormone signaling.

**SIGNIFICANCE STATEMENT:** IQD1 is involved in glucosinolate accumulation and in general defense responses. JA activates IQD1 that acts upstream or parallel to JA/ET and SA signaling pathway while controlling glucosinolate accumulation and defense against *Botrytis cinerea* and it is dependent on JAR1 controlling indole glucosinolate accumulation.

## INTRUDUCTION

Plants must continuously adapt and protect themselves both against abiotic stressors (drought, extreme temperatures, improper lighting and excessive salinity) as well as biotic stress imposed by other organisms such as viruses, bacteria, fungi and insects. Plants are resistant to most pathogens in spite of their sessile nature because they evolved a wide variety of constitutive and inducible defense mechanisms. Constitutive defenses include preformed physical barriers composed of cell walls, waxy epidermal cuticle, bark and resins (Heath, 2000a). If the first line of defense is breached, then the plant must resort to a different set of chemical mechanisms in the form of toxic secondary metabolites and antimicrobial peptides, which are ready to be released upon cell damage (Tam *et al*., 2015). These preformed compounds are either stored in their biologically active forms like saponins (Podolak *et al*., 2010), or as precursors that are converted to toxic antimicrobial molecules only after pathogen attack, exemplified by the glucosinolate - myrosinase system (Wittstock & Halkier, 2002). Other defense responses require the detection of the invading pathogen by the plant and the activation of inducible responses, often culminating in deliberate localized cell suicide in the form of the hypersensitive response (HR) in order to limit pathogen spread (Gilchrist, 1998, Heath, 2000b). Plants activate local defenses against invading pathogens within minutes and within hours, levels of resistance in distal tissue influenced by systemic signals mediated by plant hormones. The identity of the pathogen determines the type of systemic response. The classic dogma is that jasmonic acid (JA) and ethylene signaling activates resistance against necrotrophs while the salicylic acid (SA) signaling pathway is important to fight biotrophic pathogens, although it also plays some role in the defense against the necrotrophic fungi *Botrytis cinerea* (Ferrari *et al*., 2003, Govrin & Levine, 2000, Vuorinen *et al*., 2021). These two pathways are mostly antagonistic and the balance of crosstalk between them affects the outcome of the pathology (Glazebrook, 2005). *B. cinerea* causes disease in more than 200 plant species including numerous economically important crops such as tomatoes and grapes (AbuQamar *et al*., 2016). The fungus has a predominantly necrotrophic lifestyle that involves killing plant host cells by diverse phytotoxic compounds and degrading enzymes, after which it extracts nutrients from the dead cells. It comprises nearly 300 genes of Carbohydrate-Active enZymes (CAZymes) and selectively attacks the cell wall polysaccharide substrates depending on the carbohydrate composition of the invaded plant tissue (Blanco-Ulate *et al*., 2014). Plant defense response against this pathogen is complex and involves many genes related to phytohormone signaling, including the ethylene, abscisic acid, JA, and SA pathways (Kliebenstein *et al*., 2005).

Glucosinolates (GS) are sulfur rich anionic secondary metabolites characteristic of the crucifers (the Brassicaceae family) with important biological and economic roles in plant defense and human nutrition. Currently, there are approximately 140 naturally produced GS described in the literature (Nguyen *et al*., 2020). They all share a common chemical structure, consisting of a β-D-glucopyranose residue linked via a sulfur atom to a (Z)-N-hydroximinosulfate ester, plus a variable R group. GS are divided into three classes according to their precursor amino acid: compounds derived from methionine, alanine, leucine, isoleucine or valine are called aliphatic GS, those derived from phenylalanine or tyrosine are called aromatic GS and those derived from tryptophan are called indole GS. The various ecotypes of the model plant *A. thaliana* produce about 40 different GS of the indole and methionine derived aliphatic families. Glucosinolates become biologically active only in response to tissue damage, when they are enzymatically cleaved by special thioglucoside glucohydrolases known as myrosinases. These enzymes hydrolyze the glucose moiety of the GS, creating an unstable aglycone that can rearrange to form nitriles, thiocyanates, isothiocyanates and other active products. To prevent damage to the plant itself, spatial compartmentalization separates the myrosinases, which are mainly stored in specialized myrosin cells, from the GS substrates found in the vacuoles throughout the plant cells (Halkier & Gershenzon, 2006). In recent years it was demonstrated that GS metabolism is an important component of the plant defense response also against fungi and other microbial pathogens (Buxdorf *et al*., 2013, Bednarek *et al*., 2009, Clay *et al*., 2009). Regulation of GS metabolism is a complex process involving all major plant defense hormones (SA, JA, ABA and ethylene) but also other hormones such as gibberellic acid, brassinosteroids and auxin are involved (Mitreiter & Gigolashvili, 2021). Six R2R3-MYB transcription factors are known to be positive regulators of GS biosynthesis, MYB28, MYB29 and MYB76 affect aliphatic GS (Li *et al*., 2013), whereas MYB34, MYB51 and MYB122 regulate indole GS (Frerigmann & Gigolashvili, 2014, Mitreiter & Gigolashvili, 2021). IQD1 also has been found to be a positive regulator of GS accumulation and plant defense responses against insects (Levy *et al*., 2005). IQD1 is part of a family that comprises 33 IQD genes, all possessing a distinct plant specific domain of 67 conserved amino acids termed the IQ67 domain. IQ67 is characterized by a unique and repetitive arrangement of IQ, 1-5-10 and 1-8-14 calmodulin recruitment motifs (Abel *et al*., 2005). IQD genes are not unique to *A. thaliana* and bioinformatics and molecular tools have identified IQD genes in other plants such as rice, tomato, soybean, grapevine and others (Huang *et al*., 2013, Filiz *et al*., 2013, Feng *et al*., 2014, Ma *et al*., 2014, Cai *et al*., 2016, Wu *et al*., 2016, Yuan *et al*., 2019, Li *et al*., 2020). IQD genes in the plant kingdom play diverse roles unrelated to glucosinolate synthesis or defense mechanisms. A set of microarray studies directed towards identifying DELLA responsive genes identified the *A. thaliana* IQD22 as one of several proteins involved in early response to gibberllin (Zentella *et al*., 2007). The tomato IQD12 homolog SUN protein was found to be a major factor controlling the elongated fruit shape of tomato fruits (Xiao *et al*., 2008). Using virus-induced gene silencing (VIGS) method, two IQD family proteins from the cotton producing *Gossypium hirsutum* (GhIQD31 and GhIQD32), were found to induce drought and salt stress tolerance (Yang *et al*., 2019). Recent studies identified the kinesin light chain-related protein-1 (KLCR1) as an IQD1 interactor in *A. thaliana* and demonstrated association of IQD1 with microtubules. They suggest that IQD1 and related proteins provide scaffolds for facilitating cellular transport of RNA along microtubular tracks, as a mechanism to control and fine-tune gene expression and protein sorting (Abel *et al*., 2013, Bürstenbinder *et al*., 2013). The *A. thaliana* IQD16 was also implicated as being a microtubule-associated protein affecting cortical microtubules ordering, apical hook formation and cell expansion (Li et al., 2020). In the current work, we aimed to elucidate the mechanism of action of the IQD1 protein in *A. thaliana* and define its involvement in hormone signaling and in basal defense against *Botrytis cinerea*.

## RESULTS

### *IQD1* expression level correlates with *B. cinerea* resistance

Inoculation analysis with *Botrytis cinerea B. cinerea* demonstrate that the IQD1 overexpressing line (*IQD1^OXP^*) is more resistant to the necrotrophic fungus, whereas an IQD1 knockout line (*iqd1-1*) is much more sensitive (**Figure 1**).

**Figure 1.**
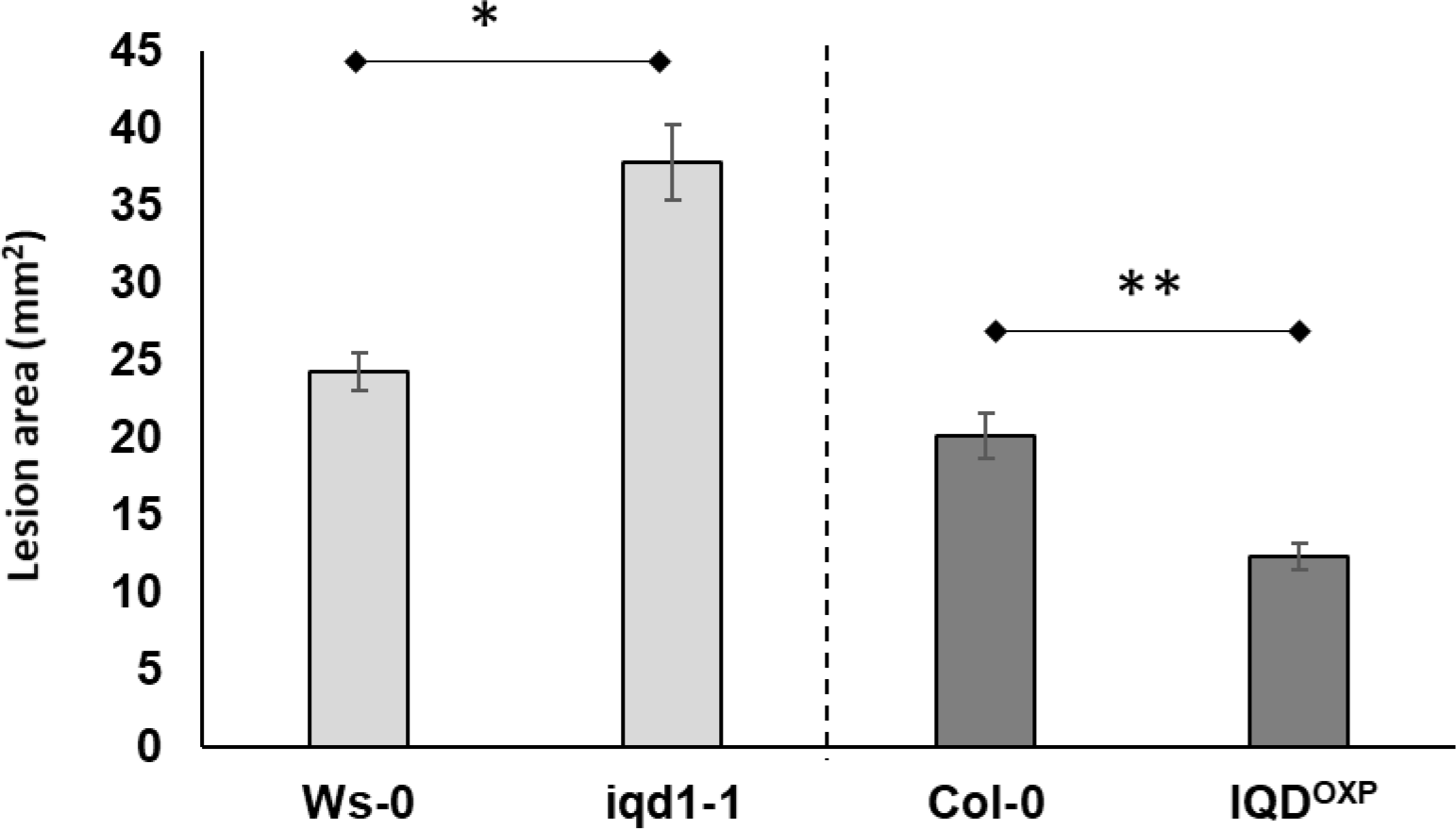
Pathogenicity of *B. cinerea* on Arabidopsis plants. Shown are averages of lesion size (mm^2^) of B. cinerea (Grape) on parental WT plants (Ws-0 or Col-0) and on IQD1 knockout line iqd1*-1* or overexpressor line IQD1^oxp^. Each column represents an average of 20 leaves with standard error bars indicated. Asterisks above the columns indicate statistically significant differences at P<0.05 from the corresponding WT, as determined using the Student t-test.

### Transcriptional characterization of the *IQD1* knockout line

#### Global gene expression analysis of iqd1-1 vs WT plants

In order to evaluate the molecular changes underlying the impact of *IQD1* expression, we performed global gene expression on RNA samples from WT and *iqd1-1* rosette leaves 48 hours after *B. cinerea* or mock inoculation.

A summary of parsed reads for each of the four samples of reads mapped to the *A. thaliana* genome is provided in **Table S1**. Our analysis revealed that 48 hours post mock inoculation, a total of 3508 genes were differentially expressed at least four-fold in *iqd1-1* knockout plants compared with WT *A. thaliana* (**Figure S1A and Table S2**). Among these genes, 1054 were upregulated in mock treated *iqd1-1* (downregulated in WT), yet more than double this number - 2454 genes exhibited downregulation in the *iqd1-1* mutant (expressed higher in WT). Eighteen genes were selected for qRT-PCR analysis in order to validate the RNA-Seq data, 7 genes that were upregulated in mock treated *iqd1-1* vs. WT and 11 genes that were downregulated in the same experiment. Expression ratios obtained by qRT-PCR were plotted versus the respective RNA-Seq values, showing that the qRT-PCR is in agreement with RNA-Seq data (**Figure S1B**).

**Table 1.**
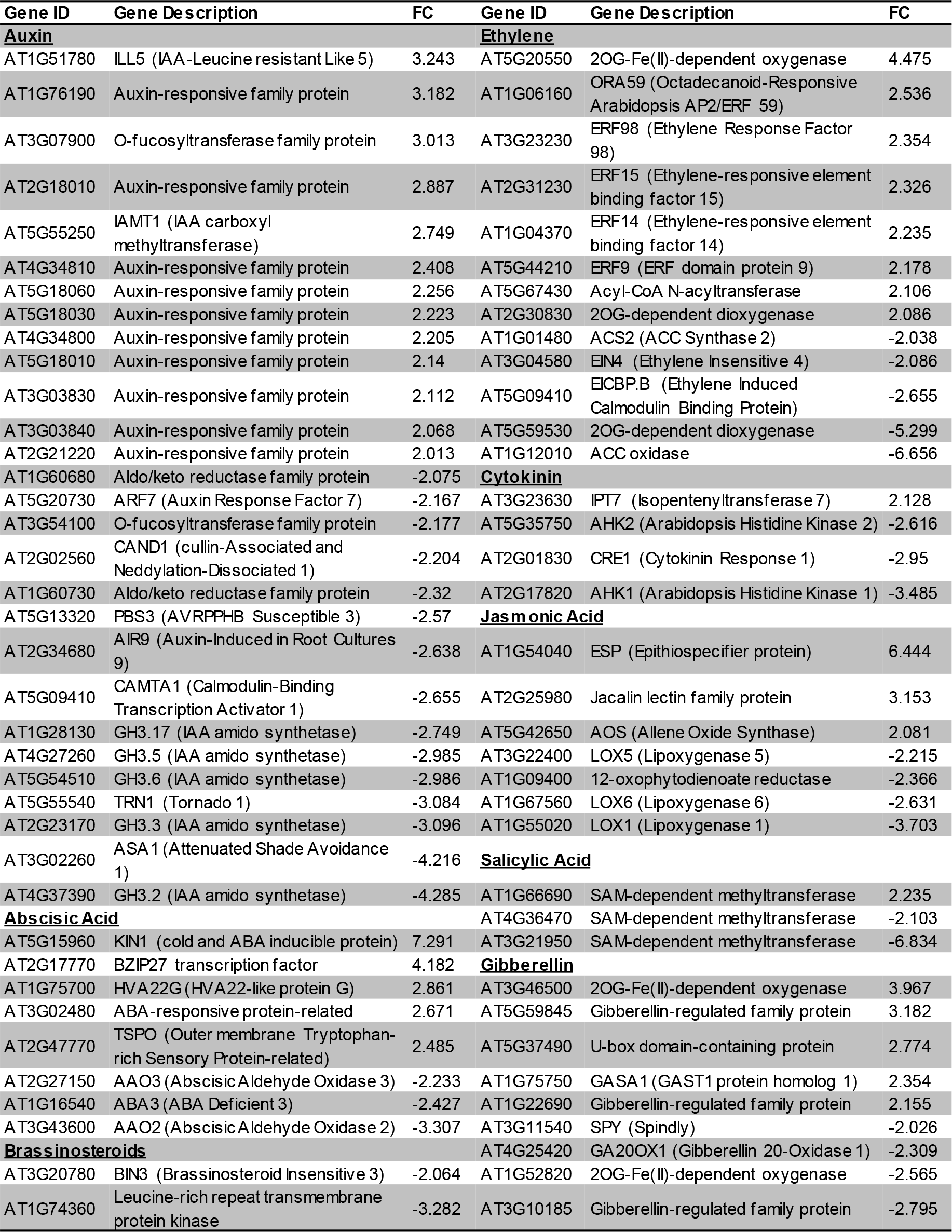
Hormone related genes differentially expressed in *iqd1-1* plants vs. WT (FC>4).

**Table 2.**
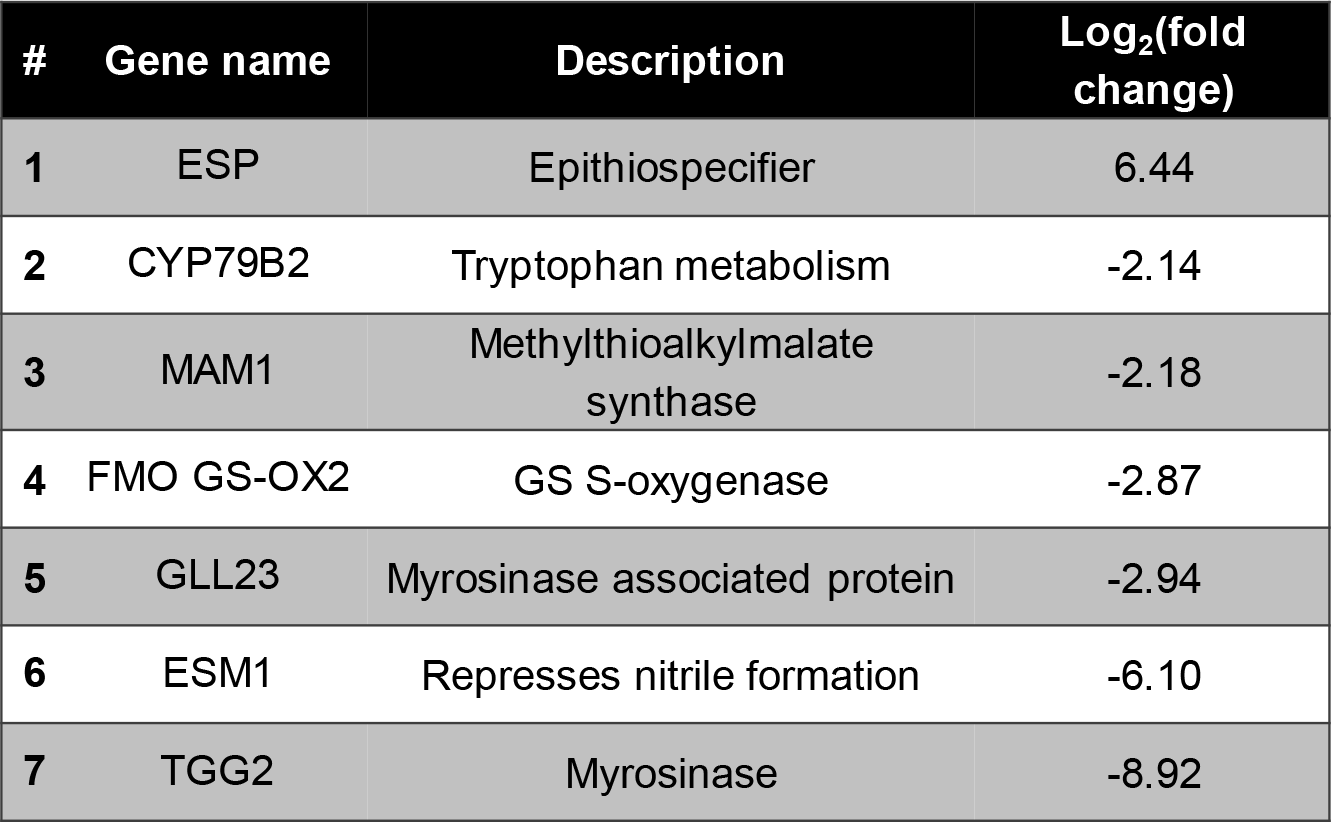
Differentially expressed genes involved in GS biosynthesis and hydrolysis in *iqd1-1* as compare to WT

| # | Gene name | Description | Log<sub>2</sub>(fold change) |
| --- | --- | --- | --- |
| 1 | ESP | Epithiospecifier | 6.44 |
| 2 | CYP79B2 | Tryptophan metabolism | -2.14 |
| 3 | MAM1 | Methylthioalkylmalate synthase | -2.18 |
| 4 | FMO GS-OX2 | GS S-oxygenase | -2.87 |
| 5 | GLL23 | Myrosinase associated protein | -2.94 |
| 6 | ESM1 | Represses nitrile formation | -6.10 |
| 7 | TGG2 | Myrosinase | -8.92 |

#### Functional annotation of DEGs

Functional annotation of our data revealed that there are many more significantly downregulated clusters in the *iqd1-1* mutant than upregulated ones. The downregulated protein families possess a wide array of functions such as molecular motors, DNA organization and repair, trans-membrane transporters, gene regulation and defense response (**Figure 2A**). It is important to notice that the second most downregulated cluster constitute the nucleotide-binding domain leucine-rich repeat (NB-LRR) plant resistance genes. These proteins are involved in the detection and initiation of specific plant defenses against diverse pathogen groups. The fact that many NB-LRR genes are expressed lower in the *iqd1-1* knockout plants, may contribute to these line’s sensitivity to pests (Levy et al., 2005). The upregulated clusters in *iqd1-1* are mainly comprised of water and lipid transporters and ethylene signaling genes, yet with lower enrichment scores than the downregulated clusters.

**Figure 2.**
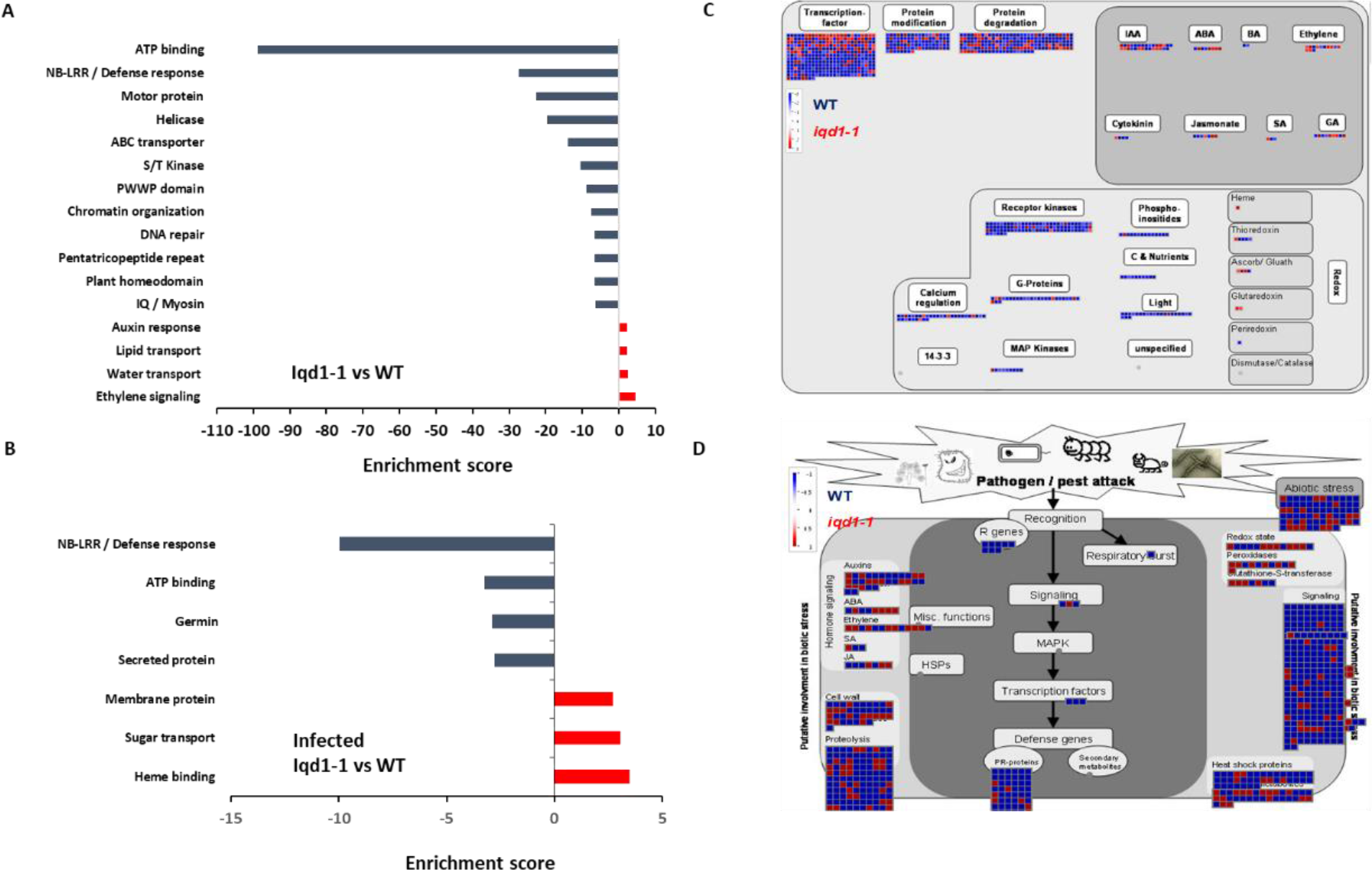
Differentially expressed clusters and genes in *iqd1-1* vs. WT plants. Enriched annotation terms of functional-related genes were grouped into clusters using the DAVID bioinformatics resources website. Positive enrichment scores denote upregulated clusters in *iqd1-1* while negative values denote upregulated clusters in WT plants. **A.** Differentially expressed clusters and genes in *iqd1-1* vs. WT plants, **B.** Differentially expressed clusters in infected *iqd1-1* vs. infected WT plants. **C-D.** MapMan regulation overview map showing differences in transcript levels between *iqd1-1* and WT. Red squares represent higher gene expression in mock treated *iqd1-1* plants while blue squares represent higher gene expression in mock treated WT plants, Regulatory network(**C**), Stress response network(**D**).

As demonstrated in Figure 2C, most of the genes assigned to plant cell regulation are downregulated in *iqd1-1* as compare to WT without any infection (**Figure 2C**, blue squares). These genes mainly function as transcription factors, protein modification and degradation, receptor kinases and hormone signaling. The only exception are ethylene-signaling genes, which are mostly upregulated in *iqd1-1* compared to WT (**Figure 2C**, red squares).

When we looked at DEGs in *iqd1-1* vs. WT that are connected to biotic stress, we found that most of the genes responsible for plant defense are downregulated in the *iqd1-1* mutant (Figure 2D, blue squares) including heat shock proteins, pathogenesis-related proteins, peroxidases and other stress response proteins. In light of the above, we can speculate that *iqd1-1* plants are impaired in sensing, signal transducing and responding to pathogen attacks. Furthermore, most of the 69 DEGs responsible for abiotic stress response are also downregulated in the *iqd1-1* mutant. They include heat shock proteins, dehydration-responsive proteins and molecular chaperones, implying impaired response to abiotic stressors as well as biotic ones. The list of depicted genes with their descriptions and fold change values are given in **Table S3**.

**Table 3.** B. cinerea differentially expressed genes with more than 50-fold change in their expression while infecting *iqd1-1* as compare to WT

| Gene annotation | Log<sub>2</sub>(FC) | Gene annotation | Log<sub>2</sub>(FC) |
| --- | --- | --- | --- |
| Cellulase | 11.96 | Hemicellulase | 6.49 |
| Extracellular membrane protein | 11.57 | MFS sugar transporter | 6.43 |
| Cellulase | 11.28 | MFS sugar transporter | 6.32 |
| Cellulase | 10.46 | MFS sugar transporter | 6.26 |
| Hemicellulase | 9.06 | Cellulase | 6.1 |
| Cellulase | 9.06 | Hemicellulase | 6.09 |
| Cellulase | 7.78 | Hypothetical protein | 6.04 |
| Transmembrane protein | 7.57 | Cellulosome complex protein | 5.95 |
| Cellulase | 7.56 | Pectinase | 5.94 |
| Hemicellulase | 7.13 | MFS transporter | 5.82 |
| Cellulase | 7.12 | Cellulase | 5.79 |
| MFS sugar transporter | 7.12 | Pectinase | 5.74 |
| MFS sugar transporter | 7.06 | Hypothetical protein | 5.71 |
| Cellulase | 6.8 | Cellulase | 5.68 |
| MFS sugar transporter | 6.7 | Pectinase | 5.67 |

### Comparison of B. cinerea infected iqd1-1 and WT plants

We found that 48 hours post inoculation with the necrotrophic fungi *B. cinerea*, 2210 genes were upregulated and 3129 genes were downregulated in infected WT compared to mock treated plants (**Figure S1A** and **Table S4**). Whereas 2343 genes were upregulated and 3092 were downregulated in infected *iqd1-1* compared to the mock treated mutant plants (Figure S1A and Table S5). Using the DAVID web resource revealed that extensive changes in gene expression occurred both in WT (**Figure S2**) and in *iqd1-1* knockout plants (**Figure S3**) after infection. In both cases, clusters that comprise gene families that participate in photosynthesis are markedly downregulated (negative values) upon infection, as the plant is tuned in to fight the invading pathogen. Upregulated clusters (positive values) consist of plant defense protein families.

Direct comparison of DEGs in infected *iqd1-1* vs. infected WT plants shows that 702 genes are upregulated in the infected mutant, while 850 genes are upregulated in infected WT plants (**Table S6**). Analysis of our RNA-Seq results revealed that WT plants express more NB-LRR resistance genes and the defensive cell-wall associated glycoproteins germins, which are induced upon pathogen recognition (**Figure 2B** negative values). On the other hand, infected *iqd1-1* plants overexpress heme-binding proteins and sugar transporters (**Figure 2B**, positive values).

### Involvement of IQD1 in hormone signaling and glucosinolate biosynthesis

#### Expression of plant hormone related genes in iqd1-1

RNA-Seq transcriptional analysis of *iqd1-1* compared to WT revealed substantial changes in gene expression in the mutant. Many of the DEGs are involved in hormone regulation and signaling (Table 1). Our analysis revealed that 35 hormone related genes were upregulated at least fourfold in *iqd1-1* plants and 37 genes were downregulated. While genes in the SA and JA pathways were mostly downregulated in *iqd1-1*, ethylene-signaling genes were noticeably upregulated. Three of the four downregulated genes in the JA pathway are lipoxygenases (*lox1*, *lox5* and *lox6*) that function as JA activated defense genes against biotic infection (Lõpez *et al*., 2011, Grebner *et al*., 2013, Viswanath *et al*., 2020). The fourth gene (*At1G09400*) is an NADPH dehydrogenase taking part in the JA biosynthesis pathway (Breithaupt *et al*., 2001). The most downregulated hormone related gene (*At3G21950*, 114.1 fold) is a salicylic acid carboxyl methyltransferase, responsible of producing a volatile methyl ester functioning as signaling molecule in systemic defense against pathogens (Chen *et al*., 2003). Five of the eight upregulated ethylene pathway genes belong to ERF/AP2 transcription factor family (*erf9*, *erf14*, *erf15*, *erf59* and *erf98*). These genes encode for ethylene response factor proteins that regulate the expression of defense responses genes following ethylene perception (Müller & Munné-Bosch, 2015). Taken together, our data indicate that IQD1 is involved in all plant hormone pathways, with strong emphasis on the major defense hormones (SA, JA and ethylene).

#### Activation of IQD1 by hormones

We observed that a large number of genes responsible for defense hormone response were altered in the *iqd1-1* line as compared to WT in the RNA-Seq results. This prompted us to investigate the effect of exogenous hormones and elicitors treatments on IQD1 expression in Arabidopsis seedlings. To this end, we used the *IQD1^pro^:GUS* reporter line that contains a fusion of the IQD1 promoter to a β-glucuronidase enzyme (Sundaresan *et al*., 1995). Histochemical staining of the reporter plants following treatment with SA or Flg22, a known activator of the SA signal transduction, showed a marked downregulation of *IQD1* expression as evident by decreased GUS staining (**Figure 3A**). In contrast, application of free JA or chitin, a major component of fungal cell walls, led to activation of *IQD1* expression, further confirming the link between *IQD1* activity and the JA pathway (**Figure 3B**).

**Figure 3.**
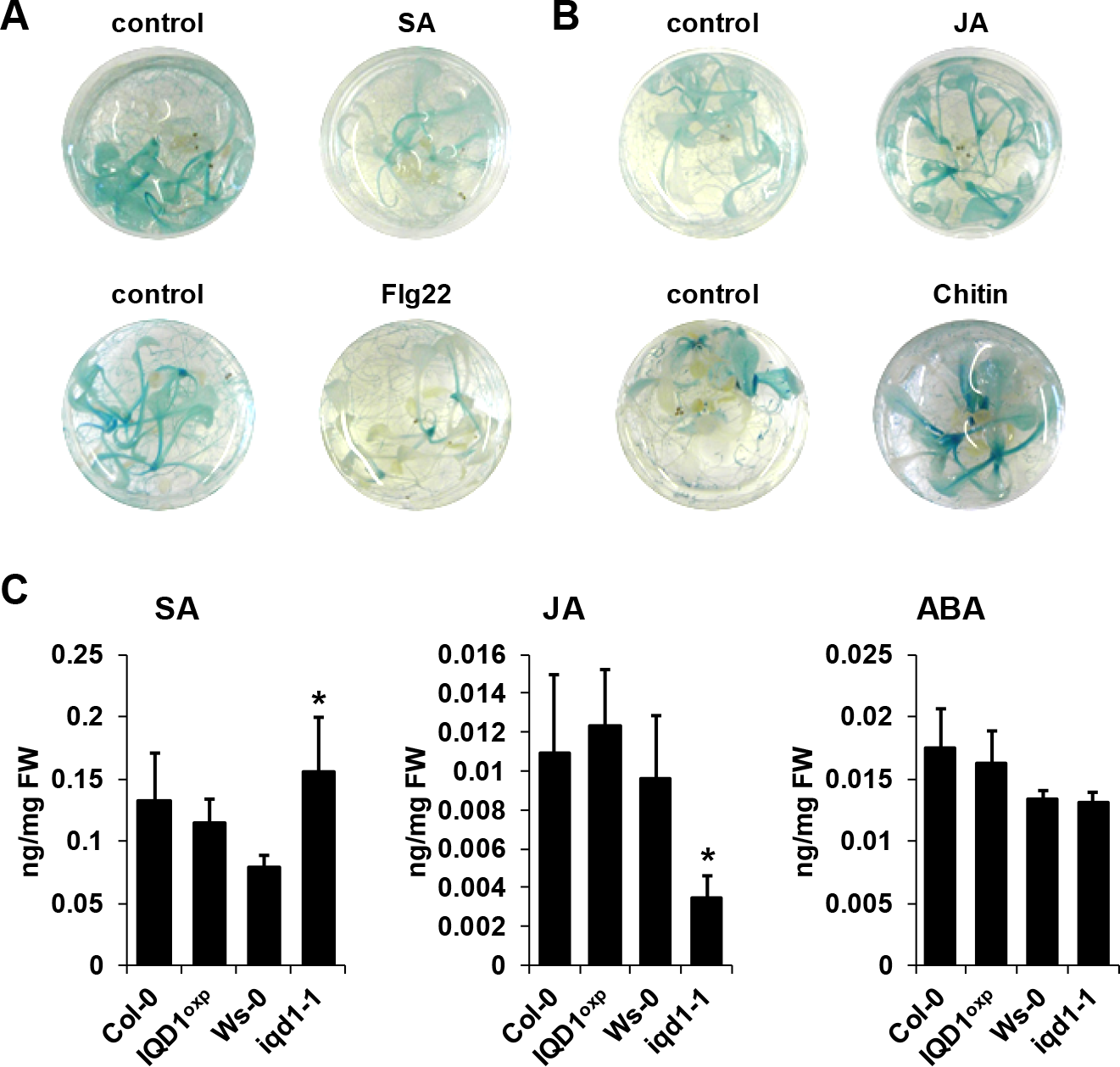
Elicitors effect on *IQD1* expression. Transgenic seedlings of gene trap line *IQD1pro:GUS* were treated with 100µM salicylic acid, 100nM Flg22 **(A)**, 100µM jasmonic acid, 500 µg/mL chitin **(B)** or an equal volume of water as control for 18 hours prior to histochemical GUS staining. (**C**) **SA, JA and ABA accumulation in *IQD1* mutants.** Plant hormones were extracted from 3 weeks old *Arabidopsis* seedlings grown on half strength MS agar plates. Quantitative analysis of plant hormones was accomplished using LC-MS/MS system and isotopically labeled analogues were used as internal standards. Each column represents an average of 3 biological replicates with standard error bars indicated. Asterisks above the columns indicate statistically significant differences compared to WT at P<0.05, as determined using student’s t-test.

We also extracted plant hormones from *iqd1-1* mutant plants and revealed significantly lower JA levels compared to WT *A. thaliana*. We also observed significantly increased SA levels but no difference in ABA levels. However, there were no changes in JA, SA or ABA content in *IQD1^OXP^* plants (Figure 3C). These results might suggest the role of IQD1 in JA accumulation and/or the synergistic effect between JA and SA signaling.

#### Dissection of IQD1 integration into defense signaling pathways

In order to investigate IQD1’s integration into defense hormone signaling pathways, we tested the epistatic relationships between *IQD1* and *A. thaliana* mutated in hormone signaling. We constructed double mutants by crossing the *IQD1^OXP^* line to mutants defective in plant-hormone signaling. Epistasis with the SA mutant *NahG* showed increased sensitivity of the single mutant compared to WT and *IQD1^OXP^*, a phenotype that was not abolished in the *NahG:IQD1^OXP^* double mutant plants (**Figure 4A**). We also determined the GS concentration in the crosses’ siblings and found that aliphatic GS content is reduced in both the single and double *NahG* mutant lines as compared to WT and *IQD1^OXP^* (Figure 4B). We observed no difference in disease severity or GS accumulation in the SA regulator mutant line *npr1* or the *npr1:IQD1^OXP^* cross (**Figure 4A, 4B**), suggesting that the dependence of GS content on SA is downstream or parallel to IQD1.

**Figure 4.**
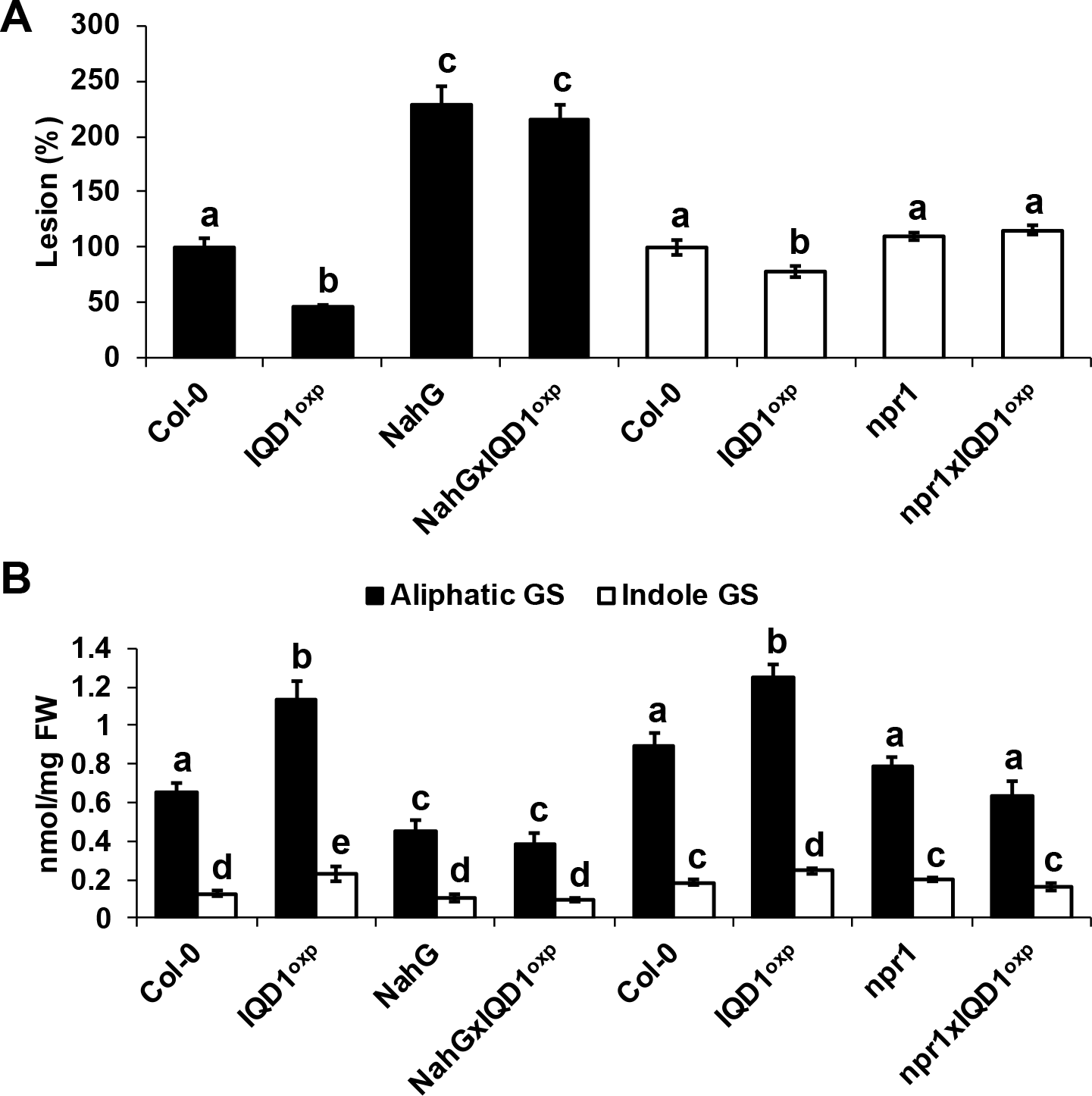
*IQD1^OXP^* effect on SA pathway mutants. **(A)** Six weeks old *Arabidopsis* detached leaves of SA pathway mutants were inoculated with *B. cinerea*. Lesion size was measured 72h post inoculation. Average lesion sizes from 30 leaves of each line are presented along with the standard error of each average. All numbers are presented as the relative percentage to their corresponding background wild-type. Different letters above the columns indicate statistically significant differences at P<0.05, as determined using the Tukey’s honest significant difference test. **(B)** Glucosinolates were extracted from six weeks old *Arabidopsis* seedlings of SA pathway mutants and analyzed by HPLC. Mean contents of methionine-derived (black bars) and tryptophan-derived (gray bars) glucosinolates are given for each line. Each column represents an average of 8 seedlings with standard error bars indicated. Different letters above the columns indicate statistically significant differences at P<0.05, as determined using the Tukey’s honest significant difference test.

All three JA signaling pathway mutant lines (*aos*, *coi1* and *jar1*) and their crosses with *IQD1^OXP^* were more resistant to *B. cinerea* infection as compared to WT. While *aos* and *aos:IQD1^OXP^* exhibited an intermediary resistance falling between *IQD1^OXP^* and WT, and *coi1* and its crossed siblings were undistinguished from *IQD1^OXP^*, *jar1* and the *jar1:IQD1^OXP^* crossed line displayed an exceptionally high resistance to *B. cinerea*, surpassing even that of *IQD1^OXP^* (**Figure 5A**). However, while GS content in *aos:IQD1^OXP^* and *coi1:IQD1^OXP^* remained unchanged compared to the parental lines, the *jar1:IQD1^OXP^* siblings displayed altered GS content. Indole GS content in the *jar1* plants was higher even than the *IQD1^OXP^* line. Indole GS concentration in the *jar1:IQD1^OXP^* siblings were lower than the *jar1* parent plants and comparable to WT levels (**Figure 5B**), suggesting that IQD1 is involved in the JA signaling pathway upstream or parallel to JAR1 and dependent on JAR1 controlling indole GS accumulation.

**Figure 5.**
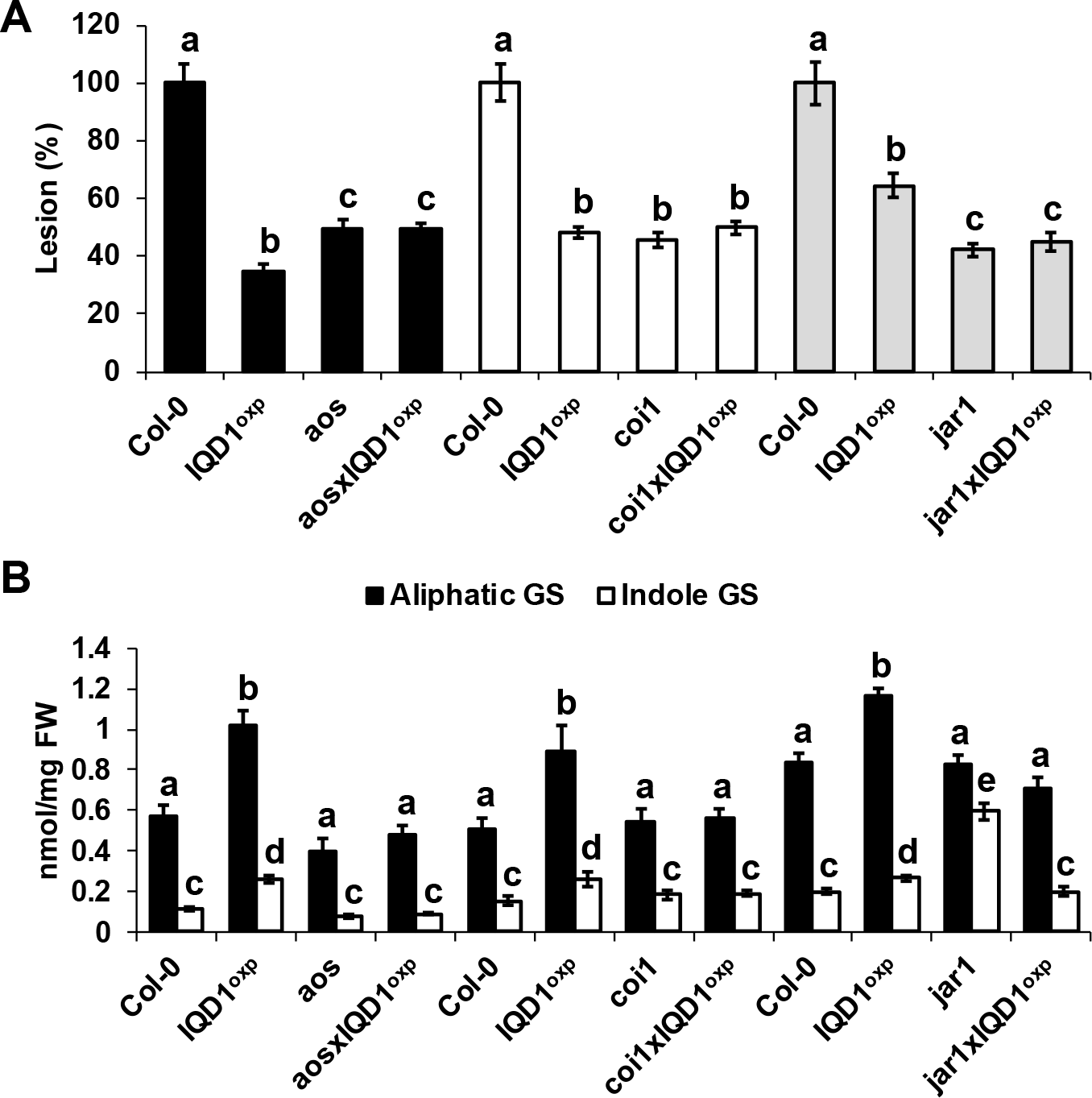
*IQD1^OXP^* effect on JA pathway mutants. **(A)** Six weeks old *Arabidopsis* detached leaves of JA pathway mutants were inoculated with *B. cinerea*. Lesion size was measured 72h post inoculation. Average lesion sizes from 30 leaves of each line are presented along with the standard error of each average. All numbers are presented as the relative percentage to their corresponding background wild-type. Different letters above the columns indicate statistically significant differences at P<0.05, as determined using the Tukey’s honest significant difference test. **(B)** Glucosinolates were extracted from six weeks old *Arabidopsis* seedlings of SA pathway mutants and analyzed by HPLC. Mean contents of methionine-derived (black bars) and tryptophan-derived (gray bars) glucosinolates are given for each line. Each column represents an average of 8 seedlings with standard error bars indicated. Different letters above the columns indicate statistically significant differences at P<0.05, as determined using the Tukey’s honest significant difference test.

As demonstrated in **Figure 6A**, Arabidopsis mutants in ethylene signaling, *ein2* and *eto1*, were more sensitive to *B. cinerea* as compared both to WT and to *IQD1^OXP^*. Siblings of *ein2:IQD1^OXP^* and *eto1:IQD1^OXP^* failed to complement this phenotype. Although indole GS levels in *eto1* plants and the crossed line *eto1:IQD1^OXP^* were higher even than *IQD1^OXP^*, it did not reflect on these lines’ resistance to *B. cinerea* infection (**Figure 6B**), suggesting that IQD1 acts upstream or in parallel to the ethylene components EIN2 and ETO1.

**Figure 6.**
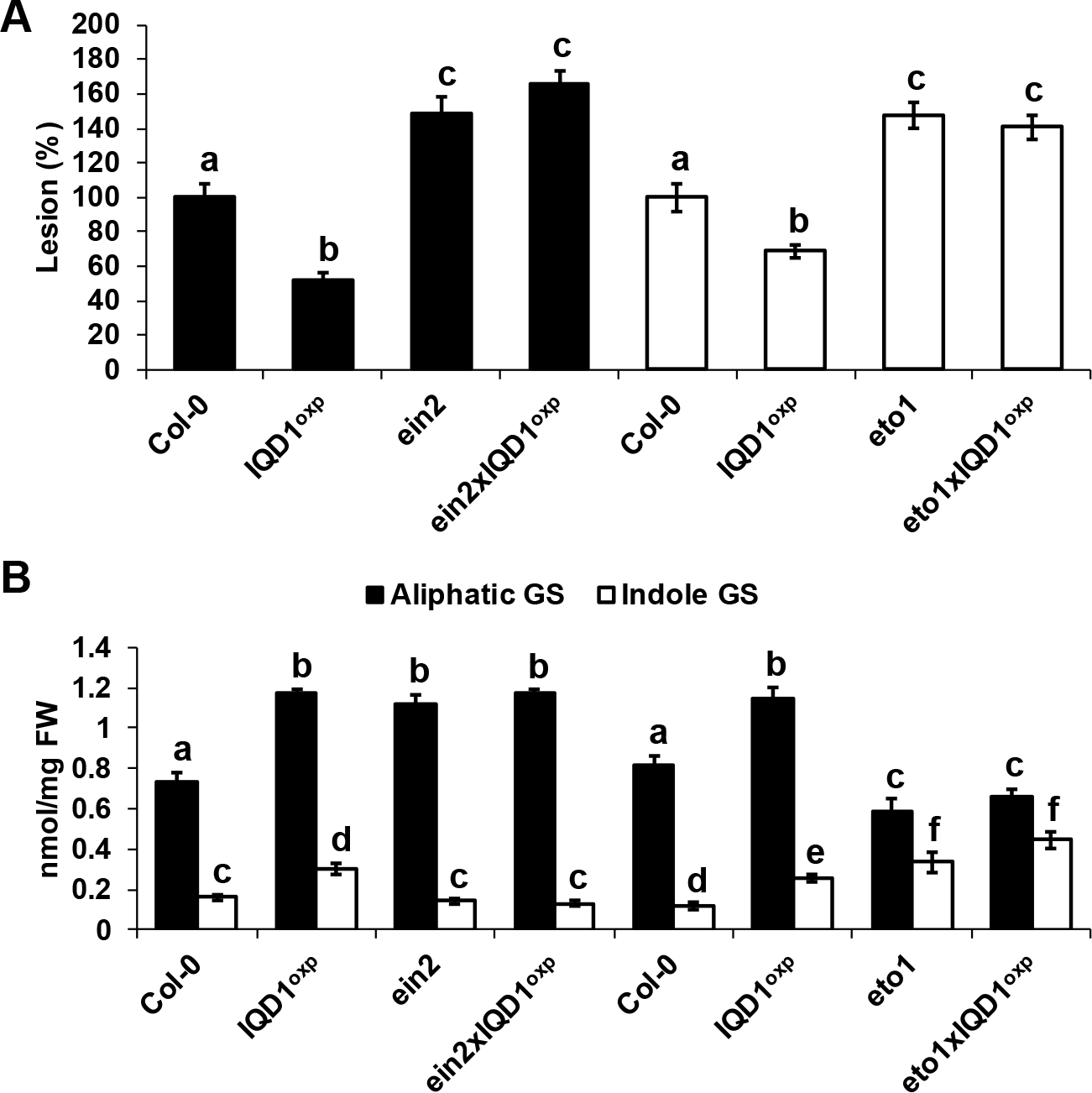
*IQD1^OXP^* effect on ethylene pathway mutants. **(A)** Six weeks old *Arabidopsis* detached leaves of ethylene pathway mutants were inoculated with *B. cinerea*. Lesion size was measured 72h post inoculation. Average lesion sizes from 30 leaves of each line are presented along with the standard error of each average. All numbers are presented as the relative percentage to their corresponding background wild-type. Different letters above the columns indicate statistically significant differences at P<0.05, as determined using the Tukey’s honest significant difference test. **(B)** Glucosinolates were extracted from six weeks old *Arabidopsis* seedlings of SA pathway mutants and analyzed by HPLC. Mean contents of methionine-derived (black bars) and tryptophan-derived (gray bars) glucosinolates are given for each line. Each column represents an average of 8 seedlings with standard error bars indicated. Different letters above the columns indicate statistically significant differences at P<0.05, as determined using the Tukey’s honest significant difference test.

#### Involvement of IQD1 in GS biosynthesis

RNA-Seq transcriptional analysis of *iqd1-1* compared to WT revealed altered expression of GS related genes. Our analysis shows that out of seven DEGs, six were downregulated in the mutant and only one was upregulated (**Figure S4**) and summarized in Table 2. Among the genes that were downregulated we find *mam1,* that encodes a methylthioalkylmalate synthase which catalyzes the condensation reactions of the first two rounds of methionine chain elongation in the biosynthesis of methionine-derived glucosinolates (Textor *et al*., 2004). The *fmo gs-ox2,* that encodes a glucosinolate S-oxygenase that catalyzes the conversion of methylthioalkyl glucosinolates to methylsulfinylalkyl glucosinolates (Li *et al*., 2008). The *cyp79b2* that belongs to the cytochrome P450 family and is involved in tryptophan metabolism (Mikkelsen *et al*., 2000). *Tgg2* a myrosinase gene involved in catabolizing GS into active products (Barth & Jander, 2006), and the *gll23* a myrosinase associated protein (Jancowski *et al*., 2014) and *esm1* that represses nitrile formation and favors isothiocyanate production during glucosinolate hydrolysis (Zhang *et al*., 2006). The only upregulated GS related gene in *iqd1-1* was *esp*, an epithiospecifier protein that promotes the creation of nitriles instead of isothiocyanates during glucosinolate hydrolysis (Lambrix *et al*., 2001). These results corroborate the active role that IQD1 participates in different steps of GS biosynthesis, as seen earlier in loss- and gain-of-function *A. thaliana* lines (Levy et al., 2005).

#### Involvement of IQD1 in *Botrytis cinerea* pathogenicity

In this study, we also analyzed the gene expression profiles of *B. cinerea* infecting the *IQD1* knockout of *A. thaliana* (*iqd1-1* mutant) compared to infection of WT plants. For statistical analysis of raw data for each sample after sequencing, see **Table S7**.

#### Identification of B. cinerea DEGs following WT and iqd1-1 infection

Unique reads that perfectly matched reference genes in each library (*B. cinerea* infecting WT or *iqd1-1)* were used to generate a matrix of normalized counts and perform statistical tests to determine whether genes are differentially expressed between pairs of factors combinations. *B. cinerea* genes with less than four-fold differences either infecting WT or infecting *iqd1-1* plants were excluded from further analyses (**Table S8**). The frequency of genes with the different fold changes in expression is shown in **Figure S5A**. A total of 678 *B. cinerea* genes were differentially expressed when it was infecting the *iqd1-1* mutant compared to the WT (fold change > 4), this includes 466 upregulated genes (expressed higher when infecting the *iqd1-1* mutant, positive values on the Y-axis) and 212 downregulated genes (expressed higher when infecting the WT, negative values on the Y-axis). Of the upregulated DEGs, 391 genes (84%) have a fold change between 4-20 and 75 genes (16%) are changed over 20-fold, reaching to a near 4000-fold difference. 194 genes (92%) of the downregulated DEGs have a fold change that is lower than 10 and only 18 genes (8%) changed more than 10-fold.

To validate the RNA-Seq data, six genes were selected for qRT-PCR analysis: *Bc1G_11623* (MFS sugar transporter), *Bc1G_10358* (hypothetical protein), *Bc1G_04691* (cellulase), *Bc1G_02144* (choline dehydrogenase), *Bc1G_12885* (MFS transporter), *Bc1G_13938* (sialidase). The expression patterns of these genes obtained by qRT-PCR and RNA-Seq are similar, indicating that the results from the RNA-Seq data are indicative of the *B. cinerea* transcriptome (**Figure S5B**).

#### Functional annotation of B. cinerea DEGs after infection of iqd1-1 mutant

The Blast2Go bioinformatics software was used in order to identify the functions of genes in the annotated *B. cinerea* genome, as more than 85% of the genes were not assigned a function (Staats and van Kan, 2012). Based on the overall analysis of gene expression profiles presented we were able to find blast hits to 460 upregulated genes (98.7%) and GO (Gene Ontology) annotations to 268 genes (57.5%) for *B. cinerea* infecting iqd1-1. The proteins encoded by the DEGs are mainly located in the plasma membrane, when classified by cellular components (**Figure 7A**). When calcified by biological processes and molecular function these proteins exhibit hydrolase activity, oxidoreductase activity and trans-membrane transporter activity and they participate in carbohydrate catabolism, oxidation-reduction processes and molecule transport across the plasma membrane (**Figure 7A**). As stated above, only 212 *B. cinerea* genes displayed higher expression levels when infecting WT plants compared to the *iqd1-1* mutant. Moreover, the difference in expression (fold change) amounted to less than 20 at most. Using the Blast2Go software, we managed to find blast hits to 204 DEGs (96.2%) and GO annotations to 115 genes (54.2%). The proteins encoded by the DEGs show a propensity for nuclear localization, when classified for cellular component. Their molecular function exhibit nucleic acid binding, helicase and kinesin activity and they participate macromolecule and nucleobase biological metabolic processes and in gene expression (**Figure 7B**).

**Figure 7.**
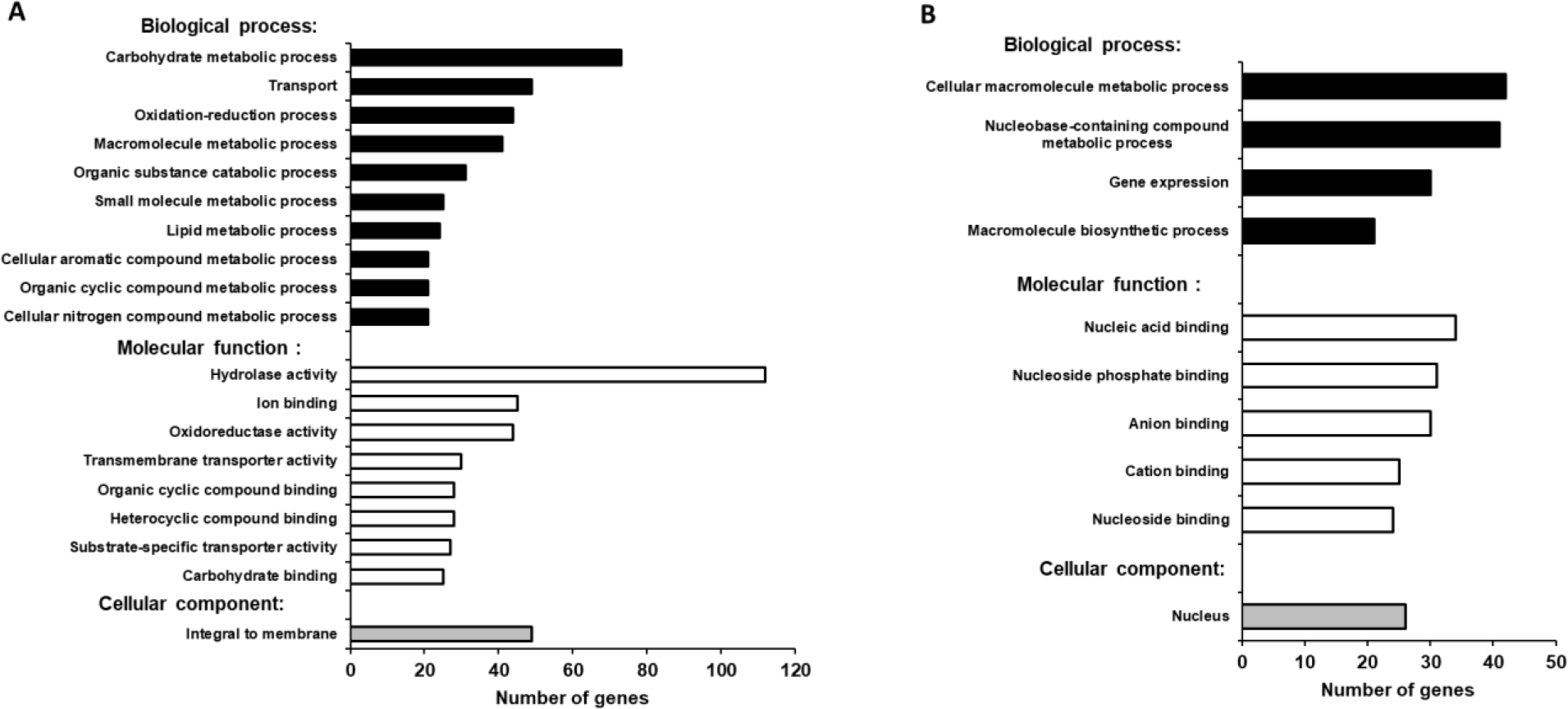
GO enrichment analysis of upregulated *B. cinerea* genes. Significantly enriched GO terms classified by biological process, molecular function and cellular component when infecting *iqd1-1* **(A)** or when infecting WT **(B).** Only GO terms applied to more than 20 differentially expressed genes are shown.

#### Highly expressed genes of B. cinerea after infection of iqd1-1 plants

To further elucidate the specific functions of the DEGs, *B. cinerea* genes with more than 50-fold changes in their expression while infecting *iqd1-1* were further analyzed. This comprised the top 30 upregulated *B. cinerea* genes while infecting *iqd1-1* (**Table 3**). The most abundant group of proteins belongs to the Carbohydrate-Active-Enzymes (CAZymes) involved in the degradation of complex carbohydrates (Garron & Henrissat, 2019). In fact, 20 out of the 30 genes (67%) in this list are CAZymes that participate in the breaking down of the host plant’s primary and secondary cell wall. These genes encode enzymes such as cellulases, hemicellulaes, pectinases and other related proteins. Seven of the DEGs (23%) belong to the Major Facilitator Superfamily (MFS) and exhibited more than 50-fold change in expression. MFS are a class of membrane proteins that facilitate the transport of small solutes such as sugars and antibiotics across the cell membrane (Yan, 2015, Niño-González *et al*., 2019). The remaining three genes in the list encode for a fungal extracellular membrane protein with anticipated role in pathogenesis, a transmembrane protein with proposed glucose transport activity and a hypothetical protein with an unknown function.

#### CAZymes distribution in DEGs

The striking number of CAZymes in the highly differentially expressed gene list, prompted us to investigate their distribution throughout the upregulated DEGs. We found that CAZymes comprise 125 out of 466 genes (27%) that are upregulated in *B. cinerea* infecting *iqd1-1*plants, while only 18 out of 212 genes (8%) were upregulated in *B. cinerea* that infect WT. The largest group (80 genes, 64%) of the CAZymes belong to the glycoside hydrolase family that constitute lytic enzymes like cellulases and hemicellulases. The second largest group (22 genes, 18%) are carbohydrate esterases that incorporate pectin catabolic enzymes. The remaining CAZymes operate on other constituents of the plant’s cell wall or play an auxiliary role to other enzymes (**Figure S6**).

## DISCUSSION

This study aimed to elucidate the molecular functions of the *A. thaliana* IQD1 protein in defense responses against the plant pathogen *B. cinerea*. A previous work with IQD1 mutants showed that the expression levels of *IQD1* in different *A. thaliana* lines is correlated with steady state accumulation of glucosinolates. Moreover, they showed that overexpressing *IQD1* has the beneficial characteristic of reducing insect herbivory of generalist insects (Levy et al., 2005). By using the necrotrophic fungal pathogen *B. cinerea*, we sought to investigate the cellular and genetic pathways in which IQD1 is regulated and affects the plant defense response. Inoculating the *IQD1^OXP^* and *iqd1-1* lines with *B. cinerea* spore suspension proved the correlation between *IQD1*’s expression levels and *A. thaliana* resistance to the fungal pathogen (**Figure 1**), as well as providing us with a simple host-pathogen system to conduct genetic screening. It was already known from our previous study that the *iqd1-1* knockout plant accumulates low levels of GS (Levy et al., 2005), In the current study, we also validate that *iqd1-1* express abnormally several of GS biosynthesis and regulation genes compared to WT plants (**Table 2**).

Information obtained from genome wide expression profiling of *iqd1-1* and WT plants following mock treatment or *B. cinerea* infection, helped us understand which plant metabolic processes were affected by the absence of IQD1. The latest genome model released for *A. thaliana* (TAIR10) contains about 27,000 protein coding genes (Lamesch *et al*., 2012). We showed that approximately 3500 genes (roughly 13% of all coding genes) were differentially expressed in the non-infected IQD1 knockout vs. WT plants (**Figure S1A** and **Table S2**). Furthermore, 70% of the genes which were downregulated in *iqd1-1* comprising diverse functions like transporters, DNA repair and gene regulation. Of notice is the large number of downregulated genes in *iqd1-1* responsible for plant defense against biotic stresses, such as cell wall remodelling proteins, signalling factors and resistance genes (**Figure 2**). Such a massive impairment of the plant defense apparatus is likely to explain the enhanced sensitivity of the knockout plants to insect and pathogen attacks (Levy et al., 2005; **Figure 1**). The ERF genes are a large family of ethylene responsive transcription factors that regulate important biological processes related to plant growth, development and plant defense (Nakano et al., 2006) (Li *et al*., 2019). This gene family was largely upregulated in the *iqd1-1* mutant (**Figure 2**, **Table 1**). The increased sensitivity to ethylene may explain several phenotypes displayed by this line such as rapid growth, large sized leaves and early development of stems and seed pods compared to WT plants (Levy et al., 2005). As demonstrated in **Figure 6** and former study, ethylene can effect glucosinolate biosynthesis (Mikkelsen *et al*., 2003) and its signalling components EIN2 and ETO1 act downstream to IQD1 controlling defense and GS accumulation.

Upon inoculation with *B. cinerea*, both the WT (**Figure S2**) and *iqd1-1* (**Figure S3**) plants have a similar basic transcriptional response. The plants shut down the energy consuming photosynthesis machinery while concentrating on fighting off the invading pathogen. The difference is that the WT plants are able to express more defense related genes like germins and R-genes (**Figure 2B**), thus resist the fungal infection more effectively than *iqd1-1*.

The three plant hormones SA, JA and ethylene play a major role in response to biotic stresses by mediating endogenous signalling that activates the expression of plant defense genes (Dong, 1998, Clarke *et al*., 2000, Li et al., 2019). Analysis of RNA-Seq data of *iqd1-1* indicates that IQD1 is involved in all three major defense hormones pathways (**Table 1**). While we see transcriptional changes in genes controlling all important plant hormones between WT and mutant plants, ethylene pathway genes are mainly upregulated in *iqd1-1* (see above), contrary to SA and JA pathway genes that show opposite characteristics (**Table 1**). Using the *IQD1^pro^:GUS* reporter line we showed that exogenous application of SA or Flg22 downregulates IQD1 expression, while JA and chitin treatment leads to the opposite effect of activating IQD1 (**Figure 3A, 3B**). Further confirmation of the link between IQD1 activity and the JA pathway came from LC/MS quantification of hormone accumulation in IQD1 mutants. We observed lower steady-state JA levels in the *iqd1-1* mutant plants compared to WT while SA levels were significantly increased (**Figure 3C**). We can speculate that IQD1 supress the accumulation of SA while activating the JA accumulation (**Figure 8**, **Figure 3C** and **Table 1**). It is clear from former publications that glucosinolate accumulation and metabolism is under control of different hormone signaling and several studies demonstrated similarly to us that changes in glucosinolate levels altered hormone levels such as JA, ET and ABA (Mikkelsen et al., 2003, Dombrecht *et al*., 2007, Malitsky *et al*., 2008, Morant *et al*., 2010, Chen *et al*., 2011, Mitreiter & Gigolashvili, 2021). As a conclusion to the results presented above, we hypothesize that the opposite effect on SA and JA levels might suggest the involvement of IQD1 in the synergistic effect between the JA and SA pathways that was well documented (**Figure 8**) (Pieterse *et al*., 2012, Koornneef & Pieterse, 2008, Li et al., 2019).

**Figure 8.**
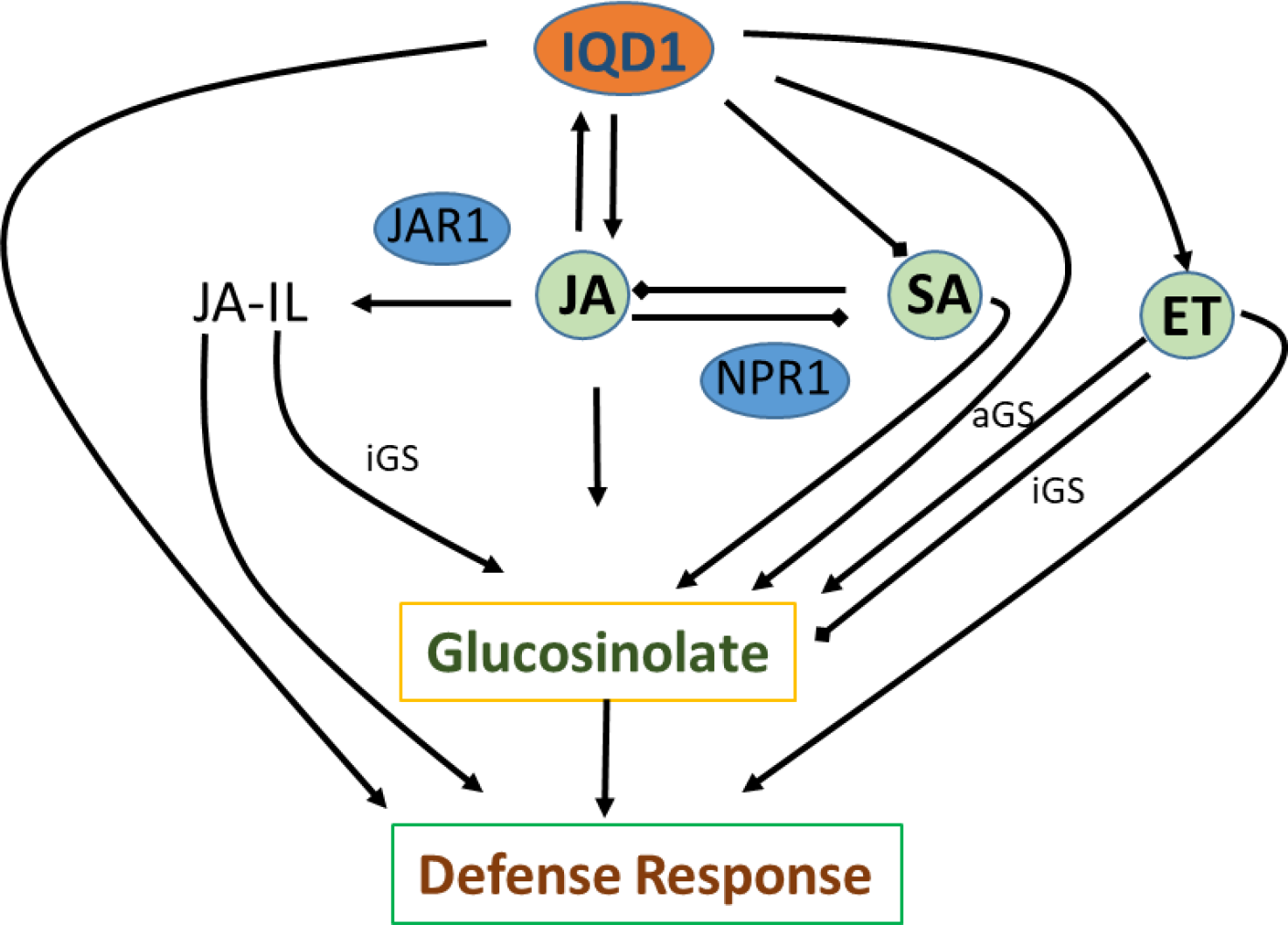
Suggested model of IQD1 involvement in glucosinolate accumulation and defense response. JA-Il, Jasmonic acid isoleucine; aGS, aliphatic glucosinolate; iGS, indolic glucosinolate.

Based on epistasis data obtained from *B. cinerea* inoculation of detached leaves and GS concentration measurement by HPLC, we managed to investigate IQD1’s integration into the three main defense hormone-signalling pathways. Overexpression of *IQD1* did not alter the resistance/sensitivity or measured GS levels in all the SA and ethylene pathway mutants we checked (**Figure 4, 6**), assuming it more likely acts parallel to them but might also act upstream for defense activation and GS accumulation (**Figure 8**). However, while indole GS content in the *jar1* plants was higher even than the *IQD1^OXP^* line, most likely due to the increase of several JA conjugates in the single mutant as described before by (Staswick and Tiryaki, 2004), the *jar1:IQD1^OXP^* cross plants accumulate significant less indole GS than the *jar1* plants and comparable to WT levels (**Figure 5**). Those results are an additional proof of the connection between IQD1 and the JA pathway. We hypothesize that IQD1 acts upstream to the JA signalling pathway and dependent on JAR1 controlling indole GS accumulation. IQD1 is also controlling JA accumulation by activating JA biosynthesis genes (**Table 1**) and activated by the JA via positive feedback loop (for model see **Figure 8**).

The extensive volume of data obtained during the RNA-Seq experiment, enabled us to investigate also the properties of *B. cinerea* infection on *iqd1-1* plants compared to WT (**Figure 7A**). Examination of the differentially expressed genes revealed that upon infection of *iqd1-1*, the fungus expresses an extensive array of Carbohydrate-Active-Enzymes (CAZymes) and membrane transporters, which facilitate the penetration and breakdown of plant tissues (**Table 3**, **Figure S6**). It has been proposed that *B. cinerea* is able to fine tune the expression of activated CAZymes according to the host’s cell wall carbohydrate composition (Blanco-Ulate et al., 2014). We hypothesize that once early penetration of the leaf tissue occurs, the fungus senses the weakness of the *iqd1-1*’s defense response and reacts by overexpressing CAZymes in order to rapidly break down the physical barriers of the plant cells (**Table 3**, **Figure S6**). We conclude that *B. cinerea* infection is more aggressive on *iqd1-1*, as the fungus takes advantage of the enhanced sensitivity of the mutant line described earlier.

In conclusion, we demonstrated in the current study that altered expression of *A. thaliana* IDQ1 has a profound effect on global expression of genes in the plant but also in the pathogen. Moreover, its expression correlates with GS levels, defense signalling and with *B. cinerea* pathogenicity function.

## EXPERIMENTAL PROCEDURES

### Plant lines and growth conditions

This work was carried out using the following *A. thaliana* (L.) Heynh. background lines: Columbia (*Col-0*), Wassilewskija (*Ws-0*) and Landsberg erecta (*Ler*). The following mutants and transgenic plants were used in *Col-0* background: *IQD1^OXP^* (Levy et al., 2005), *NahG* (Delaney et al., 1994), *npr1-1* (Cao Hui et al., 1994), *aos* (Park et al., 2002), *coi1* (Xie et al., 1998), *jar1-1* (Staswick et al., 1992), *ein2-1* (Guzmán and Ecker, 1990), *eto1-1* (Guzmán and Ecker, 1990), *pad3-1* (Zhou et al., 1999) and *cyp79B2/B3* (Zhao et al., 2002). In *Ler* background: *iqd1-2* gene trap line GT6935 (Levy et al., 2005). In *Ws-0* background: T-DNA insertion line *iqd1-1* (Levy et al., 2005). All seeds were stratified on moist soil at 4°C for 2 to 3 days before placing them in a growth chamber. Arabidopsis plants were grown at 22°C and 60% relative humidity under illumination with fluorescent and incandescent light at a photofluency rate of approximately 120 μmol m^−2^s^-1^, day length was 10 hours unless otherwise specified.

To obtain double mutants, each individual mutant was crossed with the *IQD1^OXP^* line. F1 populations were screened on Basta herbicide (glufosinate ammonium). Double homozygous mutants were identified in the F2 populations by PCR analysis with allele-specific primer pairs listed in Table S9. These plants were self-crossed and further progeny from a homozygous line was used for experiments.

### Fungal strains, growth and inoculation method

*Botrytis cinerea* (GRAPE isolate) was grown on potato dextrose agar (PDA; Difco, France) in a controlled-environment chamber kept at 22°C under fluorescent and incandescent light at a photofluency rate of approximately 120 µmol m^−2^s^−1^ and a 10/14 hours photoperiod.

Conidia were harvested in sterile distilled water and filtered through a 45 µm cell strainer to remove hyphae. For inoculation, the conidial suspension was adjusted to 1,500 conidia/µl in half-strength filtered (0.45 µm) grape juice (pure organic). Leaves were inoculated with 4 µl droplets of conidial suspension prior to RNA purification. Detached leaves from the different genotypes were layered on trays of water-agar media and inoculated with 4 µl droplets of conidial suspension. Lesions were measured using ASSESS 2.0, image analysis software for plant disease quantification (APS Press, St. Paul, MN, USA).

### GUS histochemical assay

To carry out GUS reporter gene staining assays, *iqd1-2* (GT6935 line) seeds were sterilized in (70% ethanol, 0.05% tween 20) for 5 min, washed with 100% ethanol and left to air dry. Seeds were germinated in 12-well microtiter dishes sealed with parafilm, each well containing 3 seeds and 2 ml seedling growth medium (SGM; 0.5x Murashige and Skoog basal medium with vitamins [Duchefa, Haarlem, The Netherlands] containing 0.5 g/L MES hydrate and 1% sucrose at pH 5.7). Seedlings were grown for 14 days in a growth chamber with continuous shaking at 100 rpm before treatment with elicitors. Elicitors were used at the following concentrations: 100 µM SA, 100 µM JA, 100 nM Flg22 and 500 µg/mL chitin. 18 hours after treatment with elicitors, 2 ml of GUS substrate solution (125 mM sodium phosphate pH 7, 1.25 mM EDTA, 1.25 mM K4[Fe(CN)6], 1.25 mM K3[Fe(CN)6], 0.5 mM X-Gluc and 1.25% Triton X-100) was poured in each well. The plants were vacuum-infiltrated for 10 min and then incubated at 37°C overnight covered in aluminum foil. Tissues were destained with 100% ethanol overnight and placed in 70% ethanol before digital pictures were taken.

### LC/MS quantification of salicylic, jasmonic and abscisic acid

Quantitative analysis of plant hormones was accomplished using LC-MS/MS system which consisted of a 1200 series Rapid Resolution liquid chromatography system (vacuum micro degasser G1379B, binary pump G1312B, autosampler G1367C and thermal column compartment G1316B) coupled to 6410 triple quadruple mass selective detector (Agilent Technologies, Santa Clara, CA, USA). Analytes were separated on an Acclaim C18 RSLC column (2.1×150 mm, particle size 2.2 µm, Dionex) upon HPLC conditions described in Table S10.

Mass spectrometer was operated in negative ionization mode, ion source parameters were as follows: capillary voltage 3500V, drying gas (nitrogen) temperature and flow 350°C and 10 l/min respectively, nebulizer pressure 35 psi, nitrogen (99.999%) was used as a collision gas. The LC-MS system was controlled and data were analyzed using MassHunter software (Agilent Technologies). Quantitative analysis of plant hormones was accomplished in multiple reaction monitoring (MRM) mode, isotopically labeled analogues were used as internal standards. MRM parameters are listed in Table S11.

### Glucosinolate extraction and purification

Six weeks old soil grown *A. thaliana* seedlings were weighted and lyophilized. GS were extracted with 80% methanol supplemented with sinigrin as internal standard. The extracted GS were purified on a Multiscreen 96 wells filter plate loaded with 45 µl DEAE-sephadex A25 anion exchange beads. The plate was washed once with distilled water, loaded with 200 µl of the GS extract and then washed with 80% methanol followed by two washes with distilled water. Elution was done by treating the plate with 100 µl of 3.5 mg/ml type H-1 aryl-sulfatase for an overnight reaction at room temperature, followed by a second elution with 100 µl distilled water.

#### 1.1 Glucosinolates quantification

20 µl of GS solution were run on a Thermo Scientific HPLC system at 1 ml/min. The column was a Luna C18(2), 150x4.6 mm, 5 µm (Phenomenex). The mobile phases were water (A) and acetonitrile (B), running time: 40 min. The gradient changed as follows: 1.5% B for 2.5 min, 20% B for 9 min, 20% B for 6 min, 95% B for 3 min and 1.5% B for 3 min. Afterwards, the column was equilibrated at 1.5% B for 16.5 min. The GS were detected with a UV detector at 226 nm. The amount of each GS was back calculated and expressed in nanomoles per milligram (nmols/mg) of fresh weight.

### RNA isolation

Total RNA was isolated from 6-week-old soil grown Arabidopsis rosette leaves 48 hours after inoculation with *B. cinerea*, jasmonic acid treatment or half-strength grape juice as control. RNA was extracted with TRI-Reagent (Sigma-Aldrich, St. Louis, MO, USA), followed by treatment with TURBO DNA-free (Ambion, Waltham, MA, USA) to remove genomic DNA contamination. Gel electrophoresis, NanoDrop 2000 spectrophotometer (Thermo Scientific, Waltham, MA) and TapeStation Instrument (Agilent Technologies, Santa Clara, CA) were used to determine the quality and quantity of the RNA. Following extraction, the RNA was stored at -80°C for subsequent analysis.

### cDNA library construction and sequencing

RNA samples were subjected to poly-A selection in order to select for mRNA specifically, randomly fragmented and reverse transcribed to cDNA. Adaptors that contain sample-specific indexes were ligated to the fragments in order to tag each sample and size-specific magnetic beads were used for fragment size selection. Enrichment of adaptor-bound inserts was achieved by PCR amplification, thereby enabling sample quantification for loading onto the sequencer. Illumina HiSeq 2500 system (Illumina Inc., San Diego, CA, USA) was used to sequence 50bp single reads.

Raw reads from each sample were processed by removing primer and adaptor sequences. The sequences quality per base was evaluated using FastQC v0.10.1, and low quality reads (Q-value < 30) were subsequently filtered out. The clean reads were aligned with TopHat v2.0.11 software against the *A. thaliana* genome (downloaded from the Ensembl Plants website) or the *Botrytis cinerea* genome (downloaded from the Broad Institute website) as references. Three mapping attempts were done in order to determine how many mismatches should be allowed per read (1, 3 or 5 mismatches) and the mapping files with up to 3 mismatches were used. The mapped reads were assigned to genes or transcripts based on the gene annotations file using HTSeq-count v.0.6.1 with the union mode.

### Analysis of gene expression and functional annotation

The differential gene expression was calculated by generating a matrix of normalized counts using the DESeq package v1.14.0. A threshold for false discovery rate (FDR) < 0.05 and fold change (FC) > 4 were used to determine significant differences in gene expression. Genes with FC < 4 were not considered to be differentially expressed and were therefore discarded.

Functional annotation of differentially expressed genes was carried out using DAVID (Database for Annotation, Visualization and Integrated Discovery) bioinformatics resources v6.7, the MapMan bioinformatics tool v3.5.1R2 and the Blast2Go bioinformatics software v3.1.

### Quantitative reverse-transcription PCR analysis

Total RNA (1 μg) was reverse transcribed with High Capacity cDNA Reverse Transcription Kit (Applied Biosystems, Waltham, MA, USA). Quantitative reverse transcription PCR was performed with the SYBR master mix and StepOne real-time PCR machine (Applied Biosystems, Waltham, MA, USA). The thermal cycling program was as follows: 95°C for 20 seconds and 40 cycles of 95°C for 3 seconds and 60°C for 30 seconds. Relative fold change in gene expression normalized to *Atef1a* (eukaryotic translation elongation factor 1 alpha) or *Bcactin* (*Bc1G_08198*) was calculated by the comparative cycle threshold 2^-ΔΔCt^ method. Primers used in qRT-PCR analysis of *A. thaliana* are listed in Table S12 and for *B. cinerea* in Table S13.

### Statistical analysis

Student’s t test was performed when data was normally distributed and the sample variances were equal. For multiple comparisons, one-way ANOVA was performed when the equal variance test was passed. Significance was accepted at *p* < 0.05. All experiments described here are representative of at least three independent experiments with the same pattern of results.

## DATA STATEMENT

All supporting information is available from *The Plant Journal* website.

## ACCESSION NUMBERS

IQD1 At3g09710

## Acknowledgements

Funding: IS-4210-09 from the Binational Agricultural Research and Development (BARD)

## AUTHOR CONTRIBUTIONS

OB and ML designed the experiments. OB performed the majority of the experiments and analyzed the data, with assistance from ML. OB and ML wrote the article together.

## CONFLICT OF INTEREST

The authors declare no competing financial interests.

## Notes

### Competing Interest Statement

The authors have declared no competing interest.

