## Supplementary data for "IQD1 involvement in hormonal signaling and general defense responses against *Botrytis cinerea*"

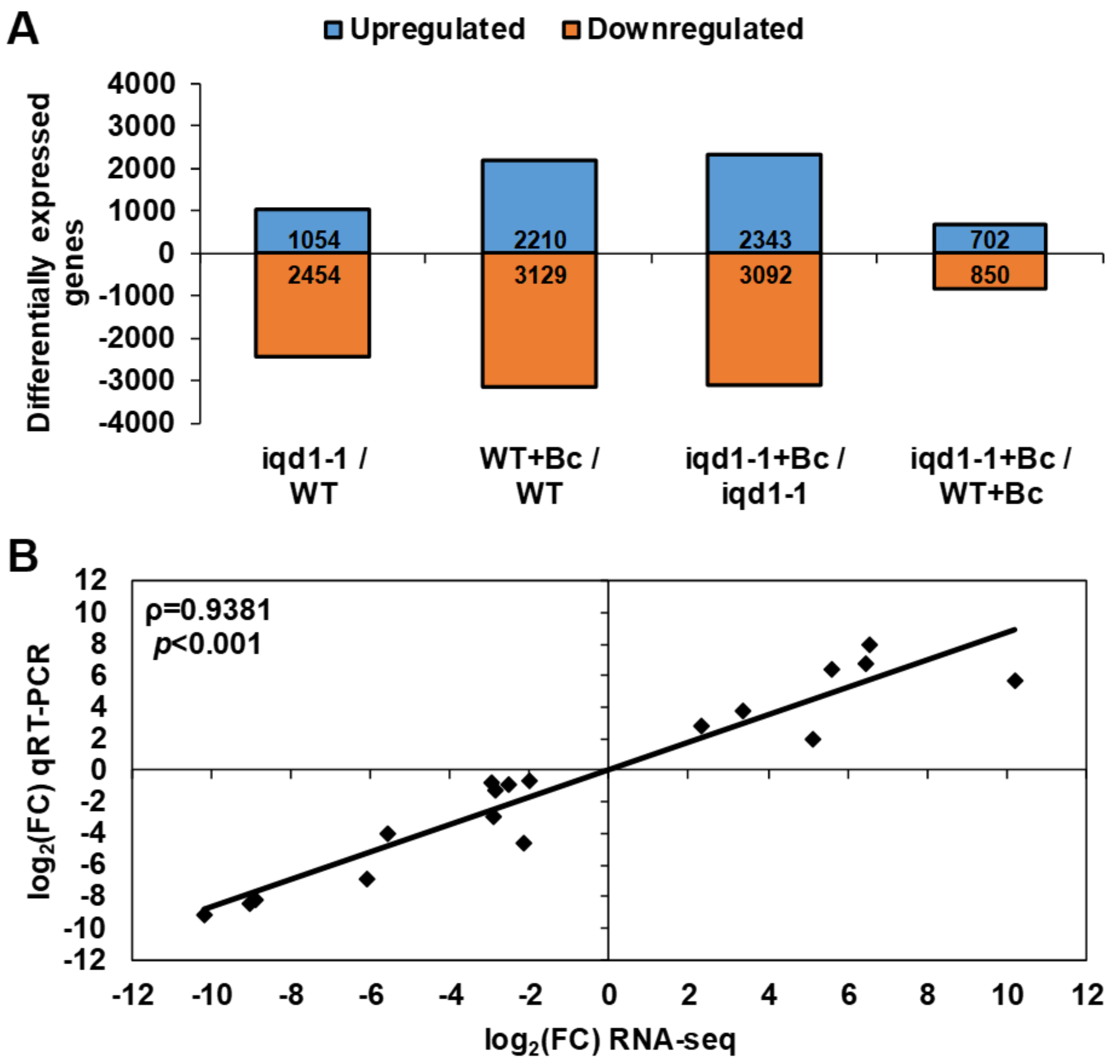


**Figure S1. A**. Differentially expressed *Arabidopsis thaliana* genes in mock and *B. cinerea* (Bc) infected plats**.** Shown are the number of upregulated and downregulated genes with more than four-fold difference between the following pairs: *iqd1-1* vs. WT, infected WT vs. WT, infected *iqd1-1* vs. *iqd1-1*, infected *iqd1-1* vs. infected WT. **B.** Correlation of qRT-PCR and RNA-Seq results. Expression ratios of 18 representative genes were determined using qRT-PCR for iqd1-1/WT. Each RNA-Seq value and qRT-PCR value (mean of 4 repeats) were $log_2$ transformed and plotted against each other for comparison. Correlation analyses were done with Spearman's Rho test.

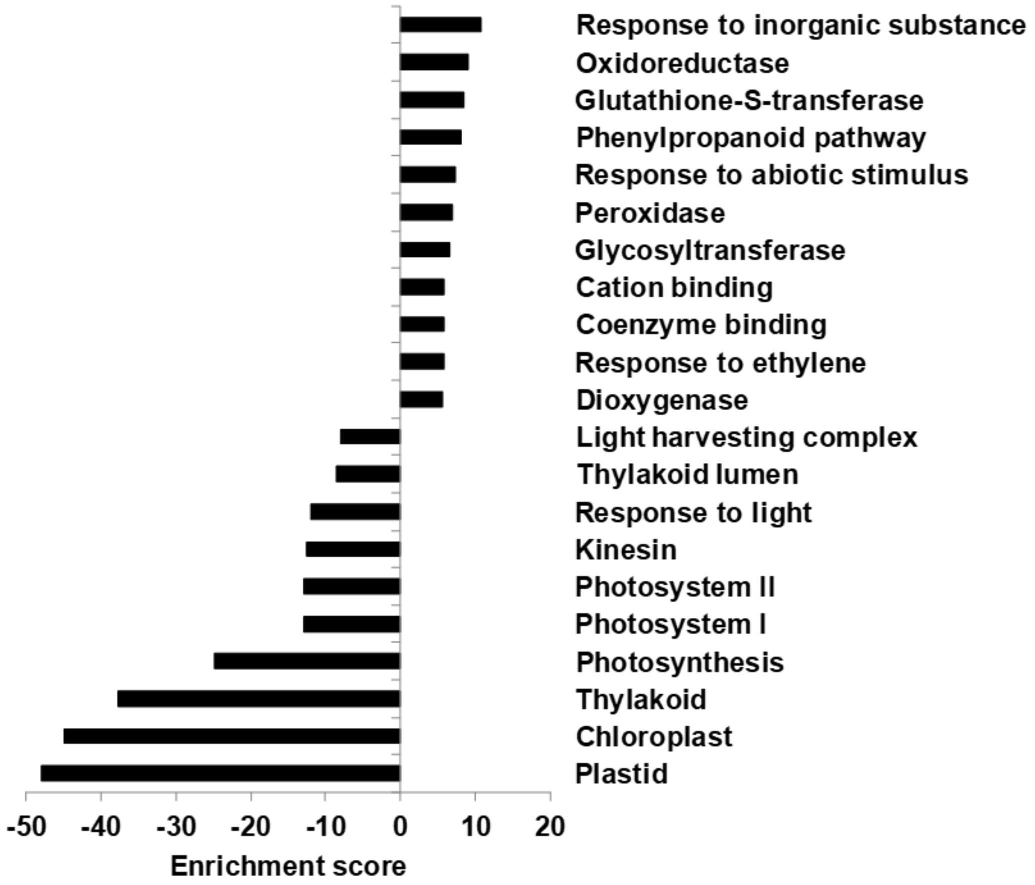


**Figure S2**. Differentially expressed clusters in infected vs. mock inoculated WT plants. Enriched annotation terms of functional-related genes were grouped into clusters using the DAVID bioinformatics resources website. Positive enrichment scores denote upregulated clusters in infected WT while negative values denote upregulated clusters in mock treated WT plants.

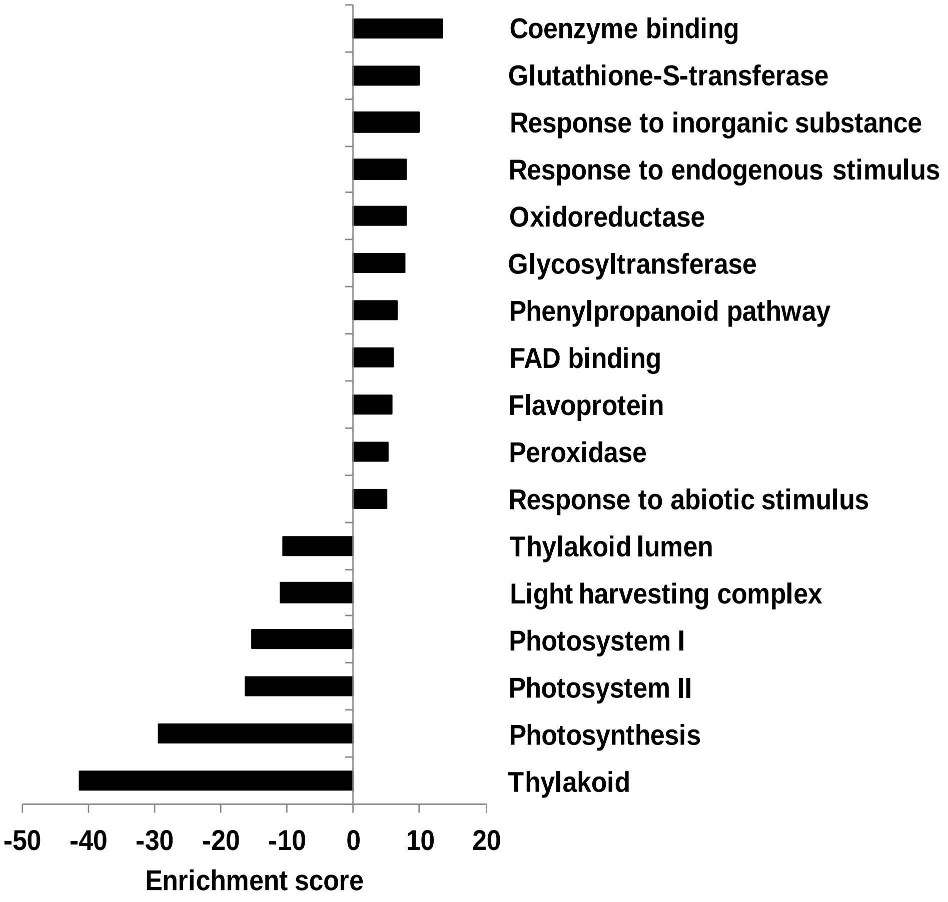


**Figure S3. Differentially expressed clusters in infected vs. mock inoculated iqd1-1 plants.** Enriched annotation terms of functional-related genes were grouped into clusters using the DAVID bioinformatics resources website. Positive enrichment scores denote upregulated clusters in infected iqd1-1 while negative values denote upregulated clusters in mock treated iqd1-1 plants.

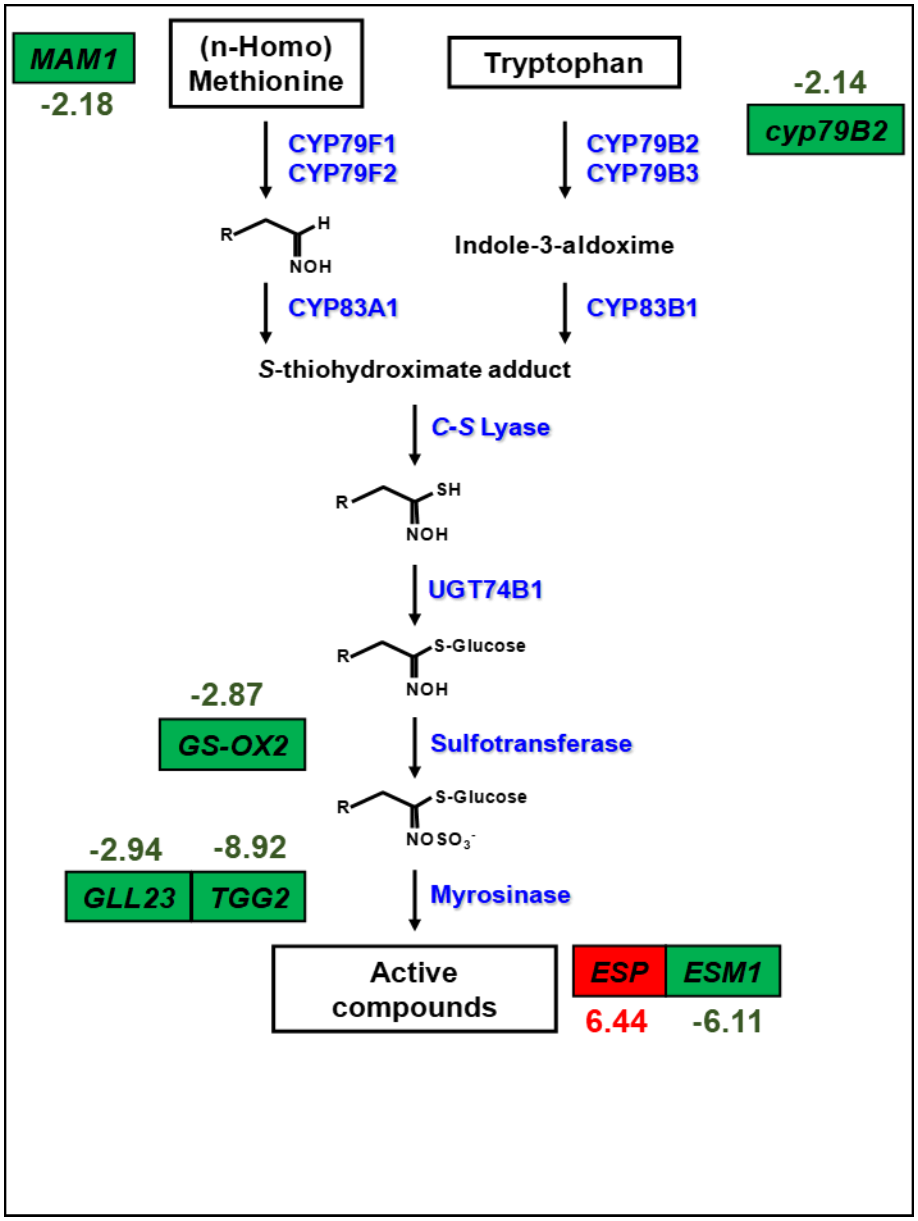


**Figure S4. Differentially expressed GS related genes in iqd1-1 compared to WT.** Schematic overview of important genes involved in glucosinolate biosynthesis. Upregulated genes in iqd1-1 are in red boxes and downregulated genes are in green boxes (numbers represent fold change).

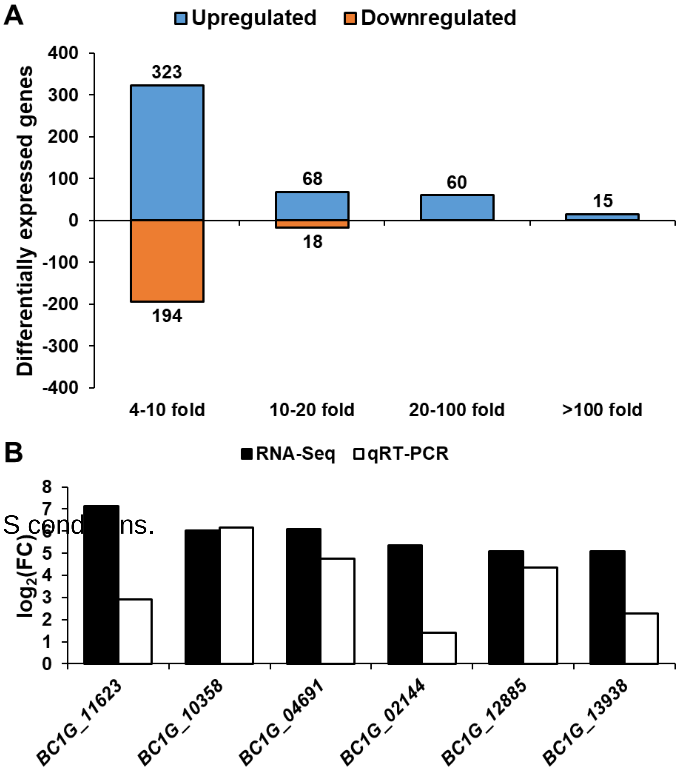


**Table S1.** LC-MS/MS cond[illegible]ns.

**Figure S5. A**. Differentially expressed Botrytis cinerea genes when infecting iqd1-1 compared to WT. Shown are the number of upregulated (positive values) and downregulated (negative values) genes with more than four-fold difference between B. cinerea infecting iqd1-1 as compared to infecting WT. **B**. qRT-PCR validation of RNA-Seq results. Comparison of B. cinerea candidate genes expression levels after infection of iqd1-1 plants vs. WT.

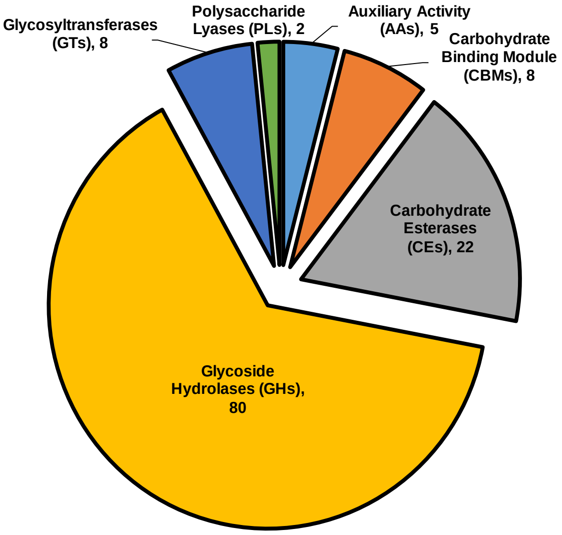


**Figure S6. Classification of CAZymes in upregulated DEGs.** The contribution of each CAZymes family is shown. Numbers in brackets denote the number of DEGs in each family.

**Table S1.** Reads of each dataset and mapping results

| Read type | Ws-0 | *iqd1-1* |
|---|---|---|
| Total reads | 45,881,751 | 47,458,716 |
| Mapped reads* | 6,359,210 (13.86%) | 3,659,067 (7.71%) |
| Uniquely mapped reads* | 3,129,135 (6.82%) | 2,477,345 (5.22%) |
| Multiple mapped reads* | 3,230,075 (7.04%) | 1,181,722 (2.49%) |
| Unmapped reads* | 39,522,541 (86.14%) | 43,799,649 (92.29%) |
| Uniquely counted reads** | 2,488,753 (79.535%) | 1,955,746 (78.945%) |
| Multiple counted reads** | 125 (0.004%) | 74 (0.003%) |
| Uncounted reads** | 640,257 (20.461%) | 521,525 (21.052%) |

**Table S7.** Summary of mapped and counted reads of RNA from *B. cinerea* and mock infected tissues.

| Read type | WT | *iqd1-1* | WT+Bc | *iqd1-1*+Bc |
|---|---|---|---|---|
| Total reads | 49,778,547 | 31,852,780 | 45,881,751 | 47,458,716 |
| Mapped reads* | 48,275,235 (96.98%) | 29,779,164 (93.49%) | 40,527,351 (88.33%) | 42,480,297 (89.51%) |
| Uniquely mapped reads* | 44,541,844 (89.48%) | 26,335,879 (82.68%) | 26,207,656 (57.12%) | 35,043,516 (73.84%) |
| Multiple mapped reads* | 3,733,391 (7.50%) | 3,443,285 (10.81%) | 14,319,695 (31.21%) | 7,436,781 (15.67%) |
| Unmapped reads* | 1,503,312 (3.02%) | 2,073,616 (6.51%) | 5,354,400 (11.67%) | 4,978,419 (10.49%) |
| Uniquely counted reads** | 43,129,103 (96.828%) | 25,238,861 (95.835%) | 24,014,417 (91.631%) | 33,296,296 (95.014%) |
| Multiple counted reads** | 636,503 (1.429%) | 433,225 (1.645%) | 357,735 (1.365%) | 556,491 (1.586%) |
| Uncounted reads** | 778,591 (1.746%) | 664,718 (2.524%) | 1,837,157 (7.010%) | 1,190,779 (3.398%) |

**Table S9.** *Arabidopsis thaliana* homozygous mutants verification primers

| Mutant | Primer name | Sequence |
|---|---|---|
| *IQD1$^{OXP}$* | H30 For | CGGAGGCGGAGAAAGTTATTG |
| | H30 Rev | CACAGGCAAAGCATTCACGAC |
| *NahG* | NahG1 f | CTGCCGCTACTCCCATATCCA |
| | NahG1 r | TCGGCTTCGGCTCGCTAC |
| *npr1-1* | npr1-1 f | AGGCACTTGACTCGGATGAT |
| | npr1-1 r | ATGCACTTGCACCTTTTTCC |
| *aos* (Salk_017756) | Salk_017756 LP | CGAGAAATTAACGGAGCTTCC |
| | Salk_017756 RP | CTAACCGGAGGCTACCGTATC |
| *coi1* (Salk_095916) | Salk_095916 LP | TCACCGACCTTCACAGATACC |
| | Salk_095916 RP | TGGTTCAAGATTGATTCCGAG |
| *jar1-1* | jar1-1 f | CAGTGTGTGTGTTTTTGATCATAAGCT |
| | jar1-1 r | CAAATTTAAACTATACCTGTTTCTGAAGG |
| *ein2-1* | ein2-1 f | CCAGAGGAAAGAGAGTTGGATGTAAAGTACTCTACCG CT |
| | ein2-1 r | CGCCATCTTTGTTTCAACAATCAGATCC |
| *eto1-1* | eto1-1 f | GCAACACAACTTGACCCTCTT |
| | eto1-1 r | GGGAGAATCCCTCAGAAAGG |
| *pad3-1* | pad3-1 f | CAAAGACATCGGGATGGCAC |
| | pad3-1 r | AGCCTTTAGCACAAGATCCACG |
| *cyp79B2* | cyp79B2 F | CTCGTTCAAGAATCCGACATCC |
| | cyp79B2 R | TCTCCGGTTTAAAGCAAAGTGG |
| *cyp79B3* | cyp79B3 F | AAGGCAATCCACCAATATCCG |
| | cyp79B3 R | TCGTGGTCAACATGCTTTATGC |

**Table S10.** LC-MS/MS conditions

| Time (minutes) | Solvent A (%), Water + 0.1% acetic acid | Solvent B (%), Methanol |
|---|---|---|
| 0 | 80 | 20 |
| 1 | 80 | 20 |
| 12 | 4 | 96 |
| 20 | 4 | 96 |
| 20.1 | 80 | 20 |
| 25 | 80 | 20 |
| Additional parameters | | |
| Column temperature (°C) | | 40 |
| Injection volume (µl) | | 5 |
| Flow rate (ml/min) | | 0.25 |

**Table S11.** MRM parameters.

| Compound | Fragmentor voltage (V) | Collision energy (eV) | MRM transitions |
|---|---|---|---|
| Abscisic acid | 80 | 10<br>10 | 263→219<br>263→204 |
| Abscisic acid D6 | 80 | 10<br>10 | 269→225<br>269→159 |
| Jasmonic acid | 80 | 10 | 209→59 |
| Jasmonic acid D5 | 80 | 10 | 213→61 |
| Salicylic acid | 80 | 15<br>15 | 137→93<br>137→65 |
| Salicylic acid D4 | 80 | 15 | 141→97 |

**Table S12.** *Arabidopsis thaliana* qRT-PCR primers

| Gene | Primer name | Sequence |
|---|---|---|
| *ef1a* | Ef1a SYBR f1 | TGAGCACGCTCTTCTTGCTTTCA |
| | Ef1a SYBR r1 | GGTGGTGGCATCCATCTTGTTACA |
| *nsp4* | NSP4 f1 | TCAAGTTTGAGTATGTCAATGGTTCTC |
| | NSP4 r1 | CAATCTCAAACTCTTCAACTCCTAGCT |
| *esp* | ESP f1 | CTACAGGAGCGAAACCTTCC |
| | ESP r1 | GATCAGGCCATACCTCACCT |
| *cml42* | CML42 f1 | TCGGATCTCGCCGAGGCGTT |
| | CML42 r1 | ACGCGACCATCTTGATTCCGGT |
| *erf114* | ERF114 f1 | CAAGTTGCGCCTACTCATCA |
| | ERF114 r1 | TTTTGGGTCTCGGATTTCAG |
| *At4G24420* | AT4G24420 f1 | ACGCTGATGAGAAGGTGATGCT |
| | AT4G24420 r1 | GGGAGTAGGGTTGGCCTTGA |
| *tip2-3* | TIP2-3 f1 | TTCTATTGGATTGCTCAGTGTCTTG |
| | TIP2-3 r1 | GGTACGCTCTTGCCATTAGTAACA |
| *At4G22490* | AT4G22490 f1 | AGTGTTGCACTGCCCTCAATG |
| | AT4G22490 r1 | CAACGGAATCGGCCTGAA |
| *sr1* | SR1 SYBR f2 | TTTGACGCCCAGGGATAGTT |
| | SR1 SYBR r2 | GGACTGTTGATAGTTTTGCTTTTGTC |
| *iqd24* | IQD24 f1 | TCCGCTTTTCGTGGCTACTT |
| | IQD24 r1 | CCCTTCACCAACGCTTGAA |
| *iqd31* | IQD31 f1 | TTCAAGATCATTCATTATCTGATGCA |
| | IQD31 r1 | GCCGAGCCAAGTAACCTCTAAA |
| *esm1* | ESM1 f1 | TCGTAGGATTGCGACAGG |
| | ESM1 r1 | CCTGAGCCTTCTCTGTGTTG |
| *tgg2* | TGG2 f1 | CCGCAAGGCCATCAAGGAGAAG |
| | TGG2 r1 | AATCTGACGGTGTAGCCGTTGC |
| *mam1* | MAM1 f1 | CGGCTGAAAGAGTTGGGATA |
| | MAM1 r1 | ATGCCTTCAAATCAGCATCC |
| *fmo gs-ox2* | GS-OX2 f1 | GCCGTTGTTGTTTGCAGTGGAC |
| | GS-OX2 r1 | ATTTCCGATGACCACCACCACCTC |
| *gll23* | GLL23 f1 | CAGACTTCCTCGCTAAATTCATGA |
| | GLL23 r1 | CTCCACGTGAGACATTGACGTT |
| *At1G63880* | AT1G63880 f1 | TGCATTCTAGCCAGCTCGAGTA |
| | AT1G63880 r1 | TCCGGGAGTTGCTTCAAATT |
| *At1G66100* | AT1G66100 f1 | TGTCTTGACATTAGTGAGTCGACATG |
| | AT1G66100 r1 | TCAGTGACAGCATCACCTGTGTT |
| *abcc11* | ABCC11 f1 | TTCAAGGTTGGGCATGATCTATC |
| | ABCC11 r1 | ACCCAAGCTGGATCACCTTCT |
| *cyp705A27* | CYP705A27 f1 | CTTATTGTGGAGCTTTTCCTTGGA |
| | CYP705A27 r1 | TGGTTAATGAGTTCGGCCATT |
| *cyp71A24* | CYP71A24 f1 | ATCTTGGATGTTTTTGTGGGTGAT |
| | CYP71A24 r1 | TGGTGGCATAGTAGCTCTGTCATT |
| *cyp76C6* | CYP76C6 f1 | TCGAACACCTACTTCTCGATATGTTT |
| | CYP76C6 r1 | TAACTCTGCCATTGCCCATTC |
| *cyp82C2* | CYP82C2 f1 | AATCTACCTGCCTGGCACTG |
| | CYP82C2 r1 | GAGAAATGGCCCATGTAAGG |
| *At1G66540* | AT1G66540 f1 | GATTGATCACTTGCTTTCTTTGCA |
| | AT1G66540 r1 | CGCAAGTATAAGAGAAAGCATGGTT |
| *cyp81K2* | CYP81K2 f1 | GTCGCTTGCTGTTGGAGCAT |
| | CYP81K2 r1 | ACGCAGGCCTCATGTCTATGT |
| *cyp76C5* | CYP76C5 f1 | CACCTACTTCTCGATATGTTTCTAGCA |
| | CYP76C5 r1 | CTCTGTCATTGCCCATTCCA |
| *cyp72A11* | CYP72A11 f1 | TCAAGACTCATGGGAGGACTTTC |
| | CYP72A11 r1 | TGTGATTTGCTCAGGATCCATT |

**Table S13.** *Botrytis cinerea* qRT-PCR primers.

| Gene | Primer name | Sequence |
| --- | --- | --- |
| *Bcactin* (*Bc1G_08198*) | 08198 F | CCCAATCAACCCAAAGTCCAACAG |
| | 08198 R | CAAATCACGACCAGCCATGTC |
| *Bc1G_11623* | 11623 F | GTTTGCATCTACCGGATCGT |
| | 11623 R | AAACACAGCACCACCATTGA |
| *Bc1G_10358* | 10358 F | GGCGAGCAAAACGCTCTAT |
| | 10358 R | GGTGAGAGGAGGAGCTTGTG |
| *Bc1G_04691* | 04691 F | CATAACCGAAAACGGCACTT |
| | 04691 R | GGCCCACTCAAAGTTATCCA |
| *Bc1G_02144* | 02144 F | GTTGATGCCACCTGGAAACT |
| | 02144 R | GGGGAGTCAAATGCGTAGAA |
| *Bc1G_12885* | 12885 F | TTCACCAAAGTCCCTCAACC |
| | 12885 R | AACCGAGGCTACCATCATTG |
| *Bc1G_13938* | 13938 F | CTCTTTTGCACTGCTTGCTG |
| | 13938 R | TTCCAGGTCTCTCCACCATC |
